# PI3Kβ inhibition creates a ferroptosis vulnerability in PTEN-deficient cancers

**DOI:** 10.64898/2026.11.24.768953

**Authors:** Yabing Nan, Tao Jiang, Fuhui Chen, Jing Ni, Yanzhi Wang, Weihua Wang, Lewis C. Cantley, Jean J. Zhao

## Abstract

PTEN loss represents one of the most frequent oncogenic events across human cancers, with PI3Kβ emerging as a dominant driver of tumor progression and therapy resistance in PTEN-deficient tumors. Although genetic or pharmacological inhibition of PI3Kβ suppresses tumor growth, actionable synthetic lethal vulnerabilities associated with PI3Kβ inhibition remain poorly defined. Here, through integrative analysis of gene expression, CRISPR dependency, and drug sensitivity datasets from the Cancer Dependency Map, we uncover ferroptosis resistance as a hallmark of PTEN-deficient cancers. Using genetically engineered mouse models (GEMMs)-derived breast cancer cells and systematic cell death screens, we show that *Pik3cb* deletion selectively sensitizes *Pten*-deficient tumors to ferroptosis. Mechanistically, PI3Kβ promotes lipid metabolic reprogramming primarily through activation of the SREBP1-dependent de novo lipogenesis, lipid droplet formation, and sequestration of ferroptosis-sensitive polyunsaturated fatty acids, thereby limiting lipid peroxidation and ferroptotic cell death. Therapeutically, combined PI3Kβ inhibition and ferroptosis induction synergistically suppress tumor growth and prolong survival across GEMMs, human cancer cell line-derived xenografts, and patient-derived xenografts. Together, these findings uncover a previously unrecognized role of PI3Kβ in ferroptosis resistance and establish a mechanistically informed combination strategy for PTEN-deficient cancers.

## INTRODUCTION

Phosphatase and tensin homolog deleted on chromosome 10 (PTEN) loss is one of the most frequent oncogenic alterations across human cancers, driving tumor progression, metastasis, and therapy resistance ^1–4^. In PTEN-deficient tumors, the phosphoinositide 3-kinase beta (PI3Kβ) isoform has emerged as a dominant driver of malignancy by promoting proliferation, survival, and immune evasion ^5–11^. Previous studies from our group and others have demonstrated that genetic or pharmacological inhibition of PI3Kβ suppresses tumor growth and enhances antitumor immunity in PTEN-deficient cancers ^4–6,8,10,12^. However, despite its critical role in PTEN-deficient tumor biology, PI3Kβ inhibition alone has shown limited durable efficacy, underscoring an incomplete understanding of therapeutic vulnerabilities in this context. Moreover, recent studies have shown that inhibition of RAS-PI3Kα interactions effectively suppresses downstream AKT signaling and exerts potent antitumor activity in KRAS-, PIK3CA-, and HER2-driven cancers, yet demonstrates minimal efficacy in PTEN-loss tumors ^13^. These findings suggest that PTEN-deficient cancers engage distinct PI3Kβ-dependent survival programs that are not effectively targeted by current PI3K-directed therapies and highlight the need to identify PI3Kβ-specific therapeutic vulnerabilities.

Ferroptosis is an iron-dependent form of regulated cell death driven by the peroxidation of polyunsaturated fatty acids (PUFAs)-containing membrane phospholipids and has emerged as a promising therapeutic vulnerability in metabolically rewired cancers ^14–23^. Multiple studies have demonstrated that specific cellular states, including mesenchymal, therapy-resistant, and drug-tolerant persister cancer cells, exhibit increased sensitivity to ferroptosis induction due to altered lipid metabolism, iron accumulation, or PUFA enrichment ^24–29^. Conversely, several oncogenic signaling pathways actively suppress ferroptosis. In particular, downstream effectors of PI3K signaling, including AKT, mTOR, and sterol regulatory element-binding protein 1 (SREBP1)-dependent lipogenic programs, have been implicated in ferroptosis resistance ^30–33^. However, the upstream PI3K isoform(s) governing ferroptosis resistance remains undefined. More importantly, whether ferroptosis represents a therapeutically exploitable vulnerability in PTEN-deficient cancers is unknown. Thus, understanding whether and how PI3Kβ-driven lipid metabolic reprogramming contributes to ferroptosis resistance in PTEN-deficient tumors may uncover novel therapeutic vulnerabilities and enable the development of more effective precision treatment strategies. Here, we investigate the role of PI3Kβ in regulating ferroptosis resistance in PTEN-deficient cancers and explore the underlying metabolic mechanisms.

## RESULTS

### PI3Kβ dependency is associated with ferroptosis resistance

Building on studies from our group and others demonstrating that PTEN-deficient cancers preferentially depend on PI3Kβ rather than PI3Kα across multiple cancer models ^5,6,8,34^, we performed integrative analyses of large-scale human cancer cell line datasets from the Cancer Dependency Map (DepMap), including gene expression, genetic dependency, and drug sensitivity profiles (**Figure S1A**). Consistent with prior observations ^35^, PTEN-deficient cancers selectively depend on PI3Kβ, but not PI3Kα, across multiple independent CRISPR and RNAi datasets (**Figure S1B**). Analysis of the Cancer Therapeutics Response Portal (CTRP) drug sensitivity dataset revealed that elevated PTEN expression or copy number variation was strongly associated with resistance to three selective PI3Kβ inhibitors, whereas no significant association was observed with the PI3Kα-selective inhibitor BYL-719 (**Figure 1A**), reinforcing PI3Kβ as a major signaling dependency in PTEN-deficient cancers.

**Figure 1.**
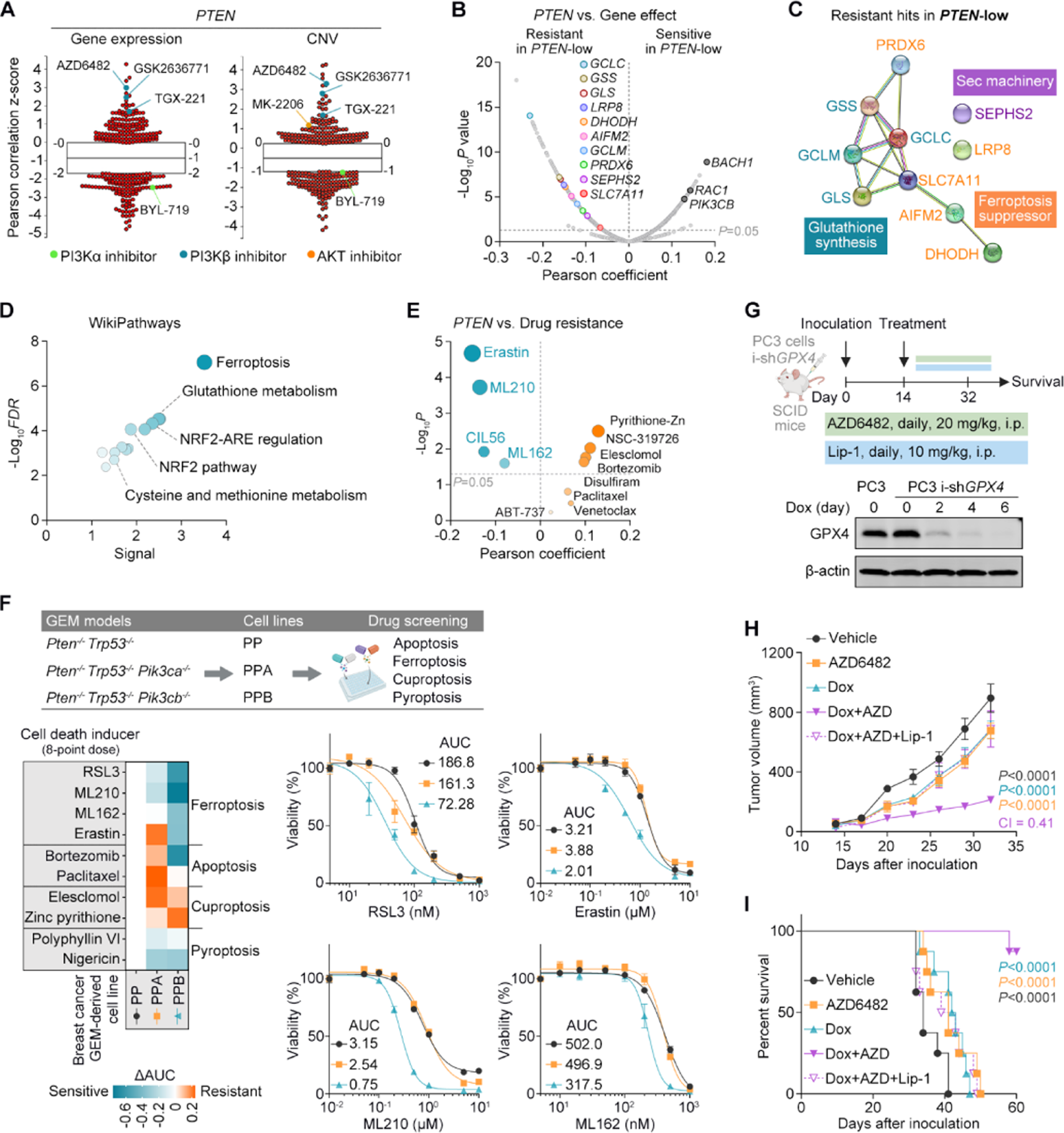
PI3Kβ inhibition synergizes with ferroptosis induction to suppress PTEN-deficient cancers. **A,** Pearson correlation analysis between PTEN expression or CNV and compound sensitivity across pan-cancer cell lines from the CTRP database. PI3K-AKT pathway inhibitors are highlighted. Values represent z-scored Pearson correlation coefficients. **B,** Pearson correlation analysis between PTEN gene expression and CRISPR dependency (gene effect) scores. **C** and **D,** STRING protein-protein interaction network and Wiki pathway enrichment analysis of the indicated selected proteins among resistant hits in PTEN-low contexts. **E,** Pearson correlation analysis between PTEN gene expression and drug resistance to the indicated cell death inducers. **F,** Schematic of the drug screen using cell death inducers in GEMM-derived primary breast cancer cells, quantification of changes in drug sensitivity (AUC), and dose-response curves for four ferroptosis inducers. Data are presented as mean ± s.d.; n = 4. **G,** Treatment scheme for PC3 xenografts with inducible GPX4 knockdown, and GPX4 protein expression in control and inducible knockdown cells. **H,** Tumor growth curves of PC3 xenografts under the indicated treatments. Data are presented as mean ± s.e.m.; two-way ANOVA; n = 8. **I,** Kaplan-Meier survival analysis of PC3 xenograft models under the indicated treatments. Kaplan-Meier method with log-rank test; n = 8.

To identify survival programs associated with PTEN loss and PI3Kβ dependency, we next analyzed correlations between PTEN expression and gene dependency scores across DepMap pan-cancer datasets. Strikingly, genes classified as top resistant hits in PTEN-low cancers were multiple canonical ferroptosis suppressors and glutathione metabolism genes, including glutamate-cysteine ligase catalytic subunit (GCLC), ferroptosis suppressor protein 1 (FSP1, encoded by apoptosis-inducing factor mitochondrial 2, *AIFM2*), solute carrier family 7 member 11 (SLC7A11), and peroxiredoxin 6 (PRDX6) (**Figure 1B**). Protein interaction network and pathway enrichment analyses revealed a strong enrichment of antioxidant and ferroptosis defense pathways among these resistant hits (**Figures 1C and 1D**). In contrast, among the top selectively essential genes included PIK3CB and Rac family small GTPase 1 (RAC1), both known mediators of PTEN-deficient tumor progression ^5,8,9,12,36^, as well as BTB and CNC homology 1, basic leucine zipper transcription factor 1 (BACH1), a transcriptional repressor that antagonizes nuclear factor erythroid 2-related factor 2 (NRF2)-mediated antioxidant function ^37^ (**Figure 1B**).

Because ferroptosis-related genes emerged prominently from these analyses, we next examined the relationship between PTEN status and sensitivity to distinct forms of regulated cell death, including apoptosis, ferroptosis, and cuproptosis. Strikingly, PTEN-low cancers were selectively resistant to multiple ferroptosis inducers, whereas no comparable association was observed for apoptosis- or cuproptosis-inducing agents (**Figure 1E**). Collectively, these findings identify ferroptosis resistance as a hallmark feature of PTEN-deficient cancers and suggest that PI3Kβ-dependent tumor fitness may, at least in part, be mediated through suppression of ferroptotic cell death.

### PI3Kβ depletion is synthetic lethal with ferroptosis induction

Given the strong association between PTEN deficiency and ferroptosis resistance, we next investigated whether suppression of PI3Kβ could create a therapeutically exploitable ferroptosis vulnerability in PTEN-deficient cancers. To identify synthetic lethal interactions associated with PI3Kβ dependency, we screened a panel of cell death-inducing compounds, targeting apoptosis, pyroptosis, ferroptosis, and cuproptosis, in genetically matched breast cancer cell lines derived from *Pten*/*Trp53*-deficient GEMMs (PP), together with additional deletion of either *Pik3ca* (PPA) or *Pik3cb* (PPB) ^8^ (**Figure 1F**). This system enabled direct comparison of PI3K isoform-specific functions within an otherwise identical PTEN-deficient genetic background. Strikingly, four independent ferroptosis inducers, RSL3, ML210, ML162, and erastin, selectively and potently induced cell death in PI3Kβ-deficient PPB cells, whereas PP and PI3Kα-deficient PPA cells remained relatively resistant (**Figure 1F**). In contrast, inducers of apoptosis, pyroptosis, and cuproptosis produced comparable responses across all genotypes (**Figures 1F and S1C**), indicating that PI3Kβ depletion selectively sensitizes PTEN-deficient cancer cells to ferroptosis rather than broadly enhancing susceptibility to all forms of cell death.

Consistently, PI3Kβ-deficient cells displayed markedly elevated lipid peroxidation following ferroptosis induction, accompanied by characteristic ferroptotic morphological changes, including extensive membrane damage and loss of cellular integrity (**Figures S1D and S1E**). Importantly, ferroptosis-specific inhibitors, including Ferrostatin-1, Liproxstatin-1 (Lip-1), and the iron chelator deferasirox, almost completely abolished PI3Kβ depletion-induced ferroptosis sensitivity, whereas inhibitors of targeting apoptosis, pyroptosis, or cuproptosis had minimal effects (**Figures S2A and S2B**). These findings demonstrate that the observed synthetic lethal interaction is highly specific to ferroptotic cell death.

To determine whether this interaction could be therapeutically exploited in vivo, we generated doxycycline-inducible glutathione peroxidase 4 (GPX4) knockdown PC3 cells, a human PTEN-deficient prostate cancer cell model, and established xenografts in mice (**Figure 1G**). Combined treatment with the selective PI3Kβ inhibitor AZD6482 (also known as KIN193) ^6^ and doxycycline-induced GPX4 knockdown significantly suppressed tumor growth and prolonged survival compared with either treatment alone, producing strong synergy with a combination index (CI) of 0.41 (**Figures 1H and 1I**). Importantly, pharmacological blockade of ferroptosis using Lip-1 largely abolished this synergistic antitumor effect of the combination (**Figures 1H and 1I**), confirming that ferroptotic cell death mediates the therapeutic interaction.

Together, these results identify ferroptosis suppression as a critical function of PI3Kβ in PTEN-deficient cancers. Genetic or pharmacological inhibition of PI3Kβ disrupts this protective state and creates a selective vulnerability to ferroptosis, establishing a synthetic lethal interaction that can be therapeutically exploited in PTEN-deficient tumors (**Figure S2C**).

### PI3Kβ suppresses ferroptosis in PTEN-deficient cancers

The findings that PTEN-deficient cancers exhibit both PI3Kβ dependency and intrinsic ferroptosis resistance prompted us to investigate whether PI3Kβ directly mediates ferroptosis suppression. Analysis of the CTRP drug sensitivity dataset revealed that resistance associated with elevated PI3Kβ expression was most strongly enriched for four canonical ferroptosis inducers, including RSL3, ML162, ML210, and erastin, whereas PIK3CA showed no comparable association (**Figure S3A**). These findings suggested a specific link between PI3Kβ signaling and ferroptosis resistance.

To further examined this relationship in PTEN-deficient cancers, we analyzed DepMap datasets and found that PI3Kβ expression positively correlated with ferroptosis pathway enrichment scores across PTEN-deficient cancer cell lines, similar to established antioxidant regulators, including NRF2 (encoded by *NFE2L2*), GPX4, and FSP1 (**Figure 2A**). We further interrogated a published genome-wide CRISPR screen performed in 786-O cells, a PTEN-deficient renal carcinoma model ^38^. Strikingly, PI3Kβ emerged as a prominent ferroptosis suppressor with dependency profiles similar to those of well-established ferroptosis suppressors, including GPX4, FSP1, and SLC7A11, whereas the other three PI3K isoforms showed little or no association with ferroptosis (**Figure 2B**).

**Figure 2.**
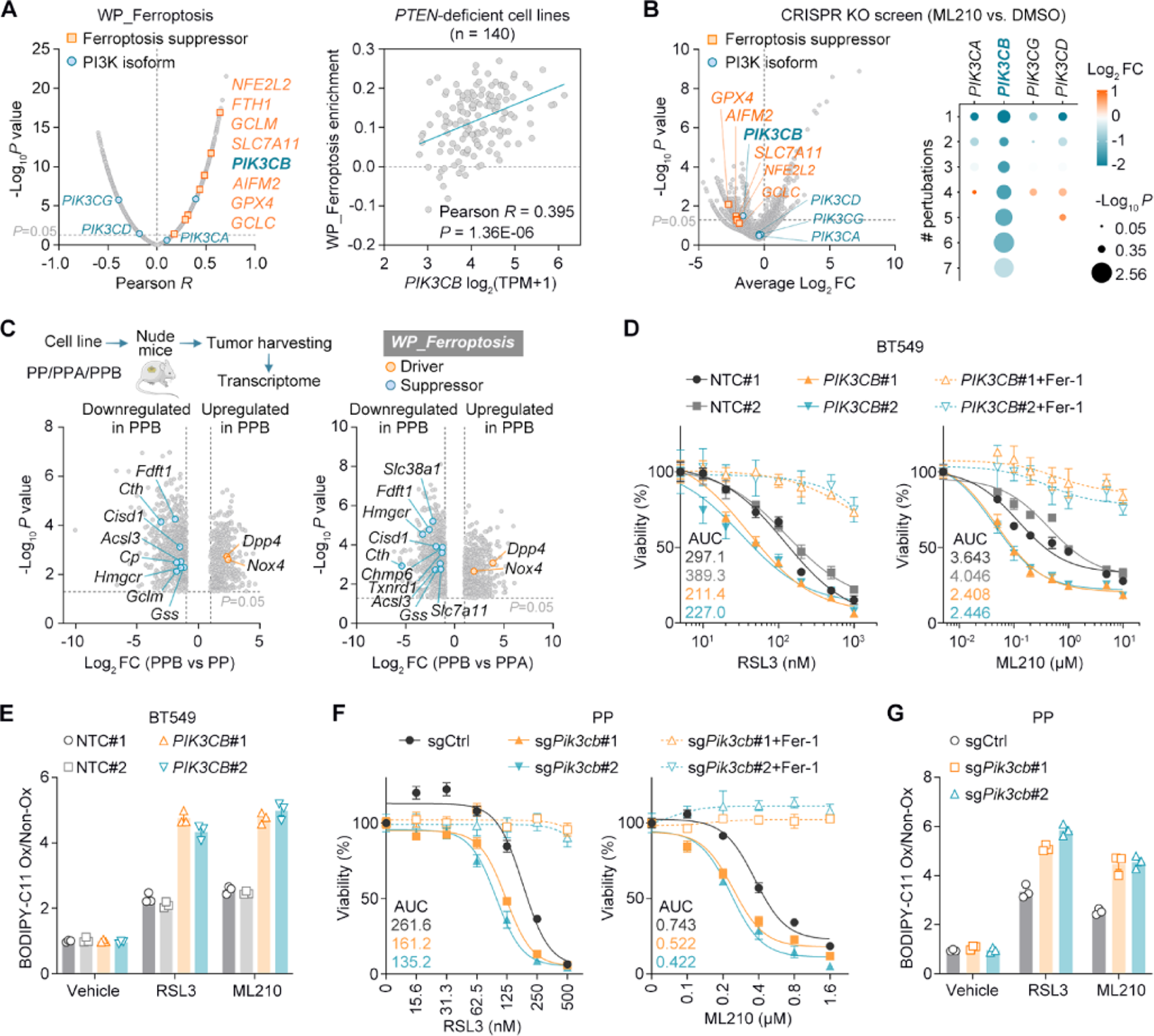
PI3Kβ inhibits ferroptosis in PTEN-deficient cancers. **A,** Pearson correlation analysis between ssGSEA ferroptosis pathway scores and gene expression in DepMap datasets. PI3K isoforms and established ferroptosis resistance genes are highlighted. **B,** Volcano plot of published CRISPR knockout screen data identifying resistance genes to the ferroptosis inducer ML210 in PTEN-deficient 786-O cells, highlighting PIK3CB as a significantly enriched resistance gene. PI3K isoforms are highlighted in green, and known ferroptosis suppressor genes are highlighted in orange. The effects of individual gRNA perturbations targeting PI3K isoforms are shown in a bubble plot. **C,** Volcano plots showing differentially expressed genes in RNA-seq data from PI3Kβ-null PPB tumors compared with PI3Kβ wild-type PP or PPA tumors. Ferroptosis resistance genes from the WP_Ferroptosis gene set are highlighted in green, and ferroptosis sensitivity genes are highlighted in orange. **D,** Dose-response curves of RSL3 and ML210 in control and CRISPRi-mediated PIK3CB-knockdown BT549 cells. Cells were pretreated with Ferrostatin-1 (Fer-1, 10 μM) for 24 h prior to treatment with RSL3 (24 h) or ML210 (48 h). Data are presented as mean ± s.d.; n = 3. **E,** Lipid peroxidation analysis in control and CRISPRi-mediated PIK3CB-knockdown BT549 cells following treatment with RSL3 or ML210. Data are presented as mean ± s.e.m.; n = 3. **F,** Dose-response curves of RSL3 and ML210 in control and CRISPR-Cas9 sgRNA-mediated Pik3cb-knockout (sgPik3cb) PP cells. Cells were pretreated with Fer-1 (10 μM) for 24 h prior to treatment with RSL3 (24 h) or ML210 (48 h). Data are presented as mean ± s.d.; n = 3. **G,** Lipid peroxidation analysis in control and CRISPR-Cas9 sgRNA-mediated Pik3cb-knockout (sgPik3cb) PP cells following treatment with RSL3 or ML210. Data are presented as mean ± s.e.m.; n = 3.

Consistently, transcriptomic analysis of GEMM-derived tumors revealed that PI3Kβ depletion led to coordinated downregulation of ferroptosis-resistance genes and upregulation of ferroptosis-promoting genes compared with either PP or PI3Kα-deficient PPA tumors (**Figures 2C and S3B**). To directly test the functional role of PI3Kβ, we depleted PI3Kβ in PTEN-deficient human cancer cell lines and GEMM-derived PP cells, as well as ectopically expressed PI3Kβ in PI3Kβ-deficient PPB cells (**Figures S3C-S3F**). PI3Kβ depletion markedly enhanced RSL3- and ML162-induced ferroptosis and lipid peroxidation in both human and murine PTEN-deficient cancer models (**Figures 2D-2G, S3G, and S3H**). Conversely, re-expression of PI3Kβ restored ferroptosis resistance and suppressed lipid peroxidation in PPB cells (**Figures S3I and S3J**). Together, these complementary computational, genetic, and functional studies identify PI3Kβ as a selective and previously unrecognized suppressor of ferroptosis in PTEN-deficient cancers.

### PI3Kβ reprograms lipid metabolism in PTEN-deficient cancers

To elucidate the mechanisms by which PI3Kβ suppresses ferroptosis, we next investigated PI3Kβ-dependent metabolic programs. Correlation analysis of PIK3CB expression against 31,117 ssGSEA pathway signatures across PTEN-deficient cancer cell lines in the DepMap database revealed that the pathways most positively associated with PIK3CB were overwhelmingly enriched for fatty acid synthesis and lipid metabolism (**Figure 3A**). Consistently, RNA-seq analysis of GEMM-derived tumors showed that PI3Kβ-deficient PPB tumors exhibited markedly suppression of fatty acid metabolism, cholesterol homeostasis, and adipogenesis compared with either wild-type PP or PI3Kα-depleted PPA tumors (**Figures 3B and S4A**). Differential expression analysis further demonstrated coordinated downregulation of genes involved in fatty acid synthesis, fatty acid activation, cholesterol biosynthesis, and lipid storage programs in PPB tumors (**Figure 3C**).

**Figure 3.**
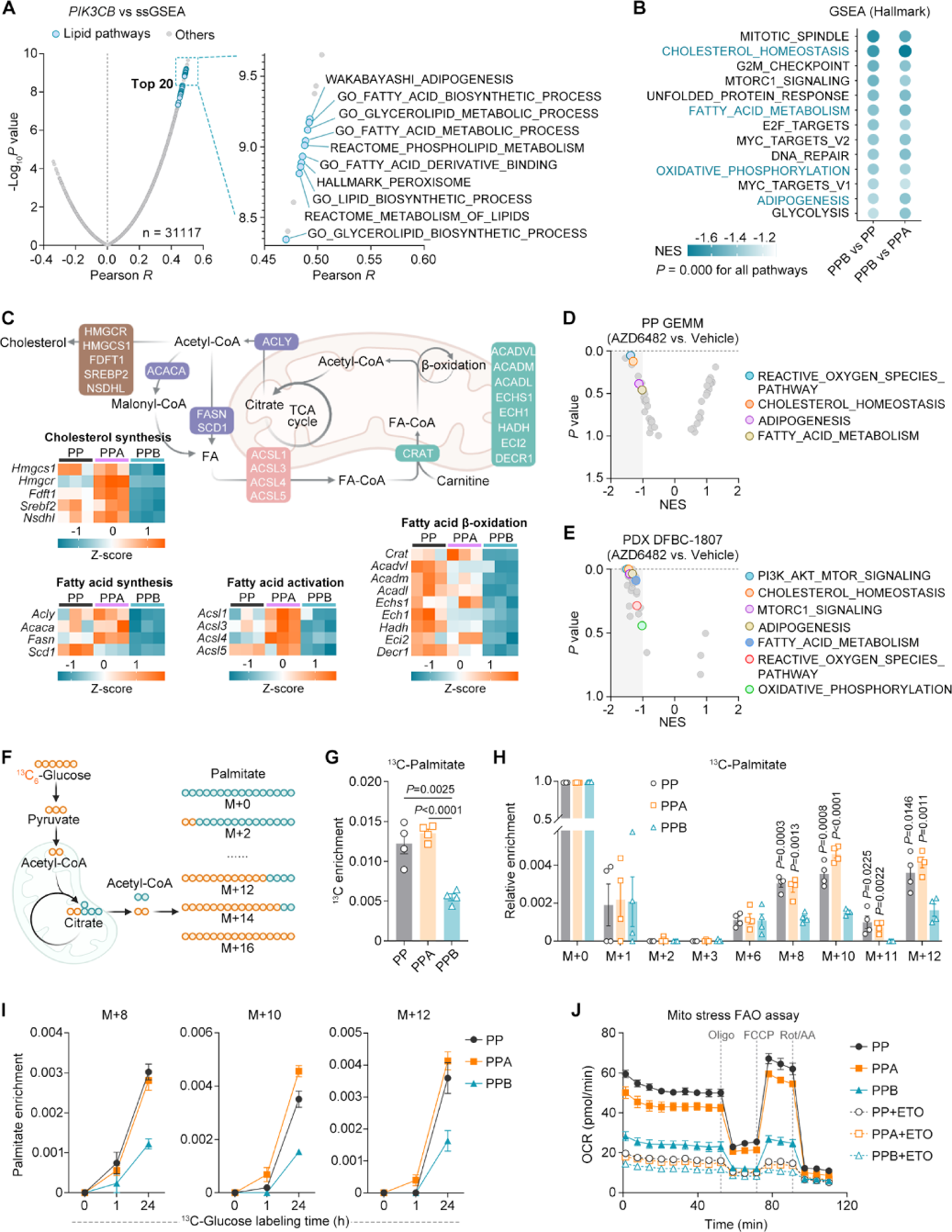
PI3Kβ reprograms lipid metabolism and promotes de novo lipogenesis in PTEN-deficient cancers. **A,** Pearson correlation analysis between PIK3CB expression and ssGSEA pathway scores in the PTEN-deficient cell lines from DepMap datasets. The top 20 significantly positively correlated pathways are shown, with 10 lipid metabolism-related pathways highlighted. **B,** Bubble plot showing GSEA enrichment results from RNA-seq data comparing PPB tumors with PP or PPA tumors. **C,** Schematic of key enzymes involved in lipid metabolism, with heatmaps showing their gene expression levels across PP, PPA, and PPB tumors based on RNA-seq data. **D,** GSEA enrichment analysis of RNA-seq data from PP tumors treated with vehicle or the PI3Kβ inhibitor AZD6482. **E,** GSEA enrichment analysis of RNA-seq data from breast cancer PDX tumors treated with vehicle or AZD6482. **F,** Schematic illustration of ^13^C_6_-glucose tracing showing isotope incorporation into the de novo lipid synthesis pathway. **G,** Total ^13^C-labeled palmitate levels in PP, PPA, and PPB cells after 24 h of ^13^C_6_-glucose labeling. Data are presented as mean ± s.e.m.; unpaired two-tailed t-test; n = 4. **H,** Fractional labeling of ^13^C-palmitate isotopologues in PP, PPA, and PPB cells following isotope tracing. Data are presented as mean ± s.e.m.; unpaired two-tailed t-test; n = 4. **I,** Time-course analysis of M+8, M+10, and M+12 palmitate isotopologues in PP, PPA, and PPB cells at 0, 1, and 24 h after isotope labeling. Data are presented as mean ± s.e.m.; n = 4. **J,** Oxygen consumption rate (OCR)-based analysis of fatty acid oxidation in PP, PPA, and PPB cells using Seahorse assays. Data are presented as mean ± s.e.m.; n = 4.

To determine whether these transcriptional programs were directly regulated by PI3Kβ signaling, we analyzed tumors treated with the PI3Kβ-selective inhibitor AZD6482. Pharmacological inhibition of PI3Kβ in PP allografts strongly suppressed fatty acid metabolism and adipogenesis pathways in vivo (**Figures 3D, S4B, and S4C**). Similar results were observed in PTEN-deficient human breast cancer patient-derived xenograft (PDX) models, where AZD6482 treatment broadly downregulated lipid metabolic programs (**Figures 3E and S4D**). Together, these findings identify PI3Kβ as a key regulator of fatty acid synthesis and lipid metabolism programs in PTEN-deficient tumors.

To functionally validate PI3Kβ-driven metabolic reprogramming, we performed untargeted metabolomics in PP, PPA and PPB cells under basal conditions and following RSL3 treatment. PI3Kβ-deficient PPB cells displayed broad downregulation of lipid-associated metabolites compared with PP and PPA cells, both before and after ferroptosis induction by RSL3 (**Figures S5A and S5B**). Pathway enrichment analysis further identified lipid metabolism and fatty acid utilization as major metabolic processes regulated by PI3Kβ signaling (**Figures S5C and S5D**).

We next directly assessed de novo lipogenesis (DNL) using ^13^C_6_-glucose tracing in PP, PPA, and PPB cells (**Figure 3F**). PI3Kβ depletion significantly reduced the incorporation of glucose-derived carbon into palmitate, indicating impaired fatty acid synthesis (**Figure 3G**). Consistently, the abundance of newly synthesized palmitate isotopologues (M+8 to M+12) was substantially diminished in PPB cells compared with PP and PPA cells (**Figure 3H**). Time-course labeling experiments further confirmed a sustained defect in DNL following PI3Kβ loss (**Figure 3I**).

Because transcriptomic and metabolomic analyses also suggested alterations in fatty acid utilization, we performed Seahorse-based fatty acid oxidation assays. PI3Kβ depletion or knockdown impaired mitochondrial respiration and fatty acid oxidation in both murine and human PTEN-deficient breast cancer cells (**Figures 3J and S5E-S5G**). Collectively, these integrated transcriptomic, metabolomic, and metabolic flux analyses demonstrate that PI3Kβ orchestrates broad lipid metabolic reprogramming in PTEN-deficient cancers, promoting DNL and lipid accumulation while reshaping fatty acid utilization. These findings raised the possibility that PI3Kβ-driven lipid metabolic programs restrict the availability of ferroptosis-sensitive PUFA, thereby contributing to ferroptosis resistance.

### PI3Kβ depletion reduces PUFA-containing neutral lipids

Given that PI3Kβ-driven DNL may serve as a key mechanism of ferroptosis defense, we performed untargeted lipidomics in PP, PPA, and PPB cells to determine the impact of PI3Kβ on lipid subclasses. Notably, PI3Kβ-mediated lipid alterations were predominantly observed in neutral lipids, particularly triacylglycerols (TAGs) and cholesteryl esters (CEs), the principal constituents of lipid droplets (**Figure 4A**). PI3Kβ depletion selectively reduced PUFA-containing TAGs and CEs, particularly those enriched in ferroptosis-sensitive arachidonic acid (AA, C20:4) and docosahexaenoic acid (DHA, C22:6) (**Figure 4B**). In contrast, neutral lipids containing ferroptosis-resistant saturated fatty acids (SFAs) or monounsaturated fatty acids (MUFAs), which are generally less susceptible to lipid peroxidation, were relatively enriched in PPB cells compared to PP and PPA cells (**Figure 4B**). Moreover, PI3Kβ-deficient PPB cells exhibited a reciprocal increase in PUFA-containing phosphatidylcholine (PC) and phosphatidylethanolamine (PE) species, lipid classes that serve as major substrates for ferroptotic lipid peroxidation, whereas MUFA- and SFA-containing PC and PE species were correspondingly reduced (**Figure 4B**).

**Figure 4.**
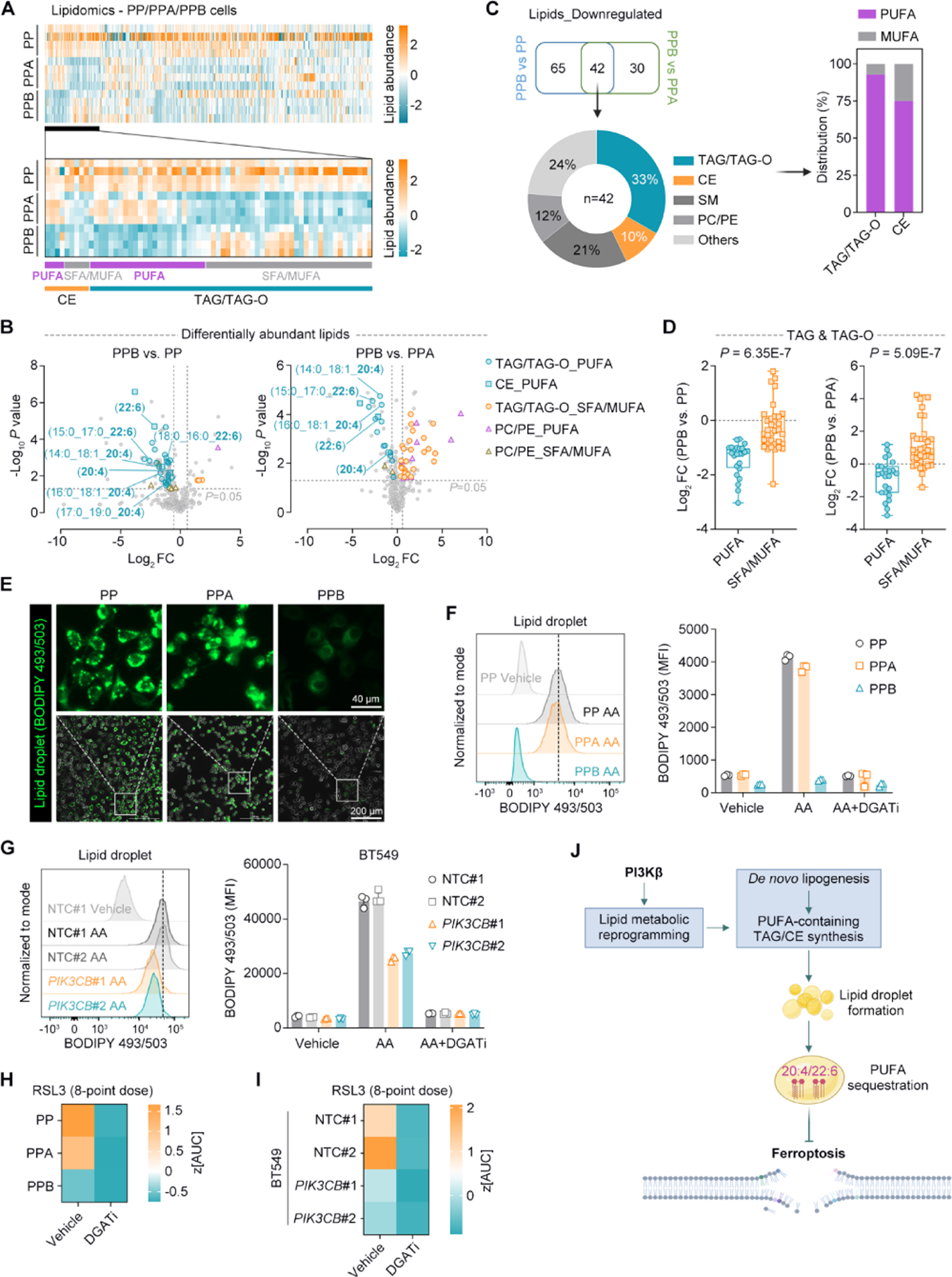
PI3Kβ promotes lipid droplet formation and PUFA sequestration to suppress ferroptosis. **A,** Heatmap showing lipid abundance from lipidomics profiling of PP, PPA, and PPB cells. **B,** Volcano plots of differentially enriched lipids in PPB compared with PP or PPA cells. PUFA-containing triglycerides (TAG) and cholesteryl esters (CE) are significantly reduced in PPB cells. **C,** Venn diagram showing 42 lipids commonly downregulated in PPB versus PP and PPB versus PPA. A pie chart illustrates that 43% of the downregulated lipids are TAG/TAG-O and CE, with bar plots further showing the distribution of PUFA- and MUFA-containing species within TAG/TAG-O and CE pools. **D,** Fold-change analysis of PUFA- and SFA/MUFA-containing TAG/TAG-O species in PPB versus PP and PPB versus PPA. **E,** Representative images of lipid droplets stained with BODIPY 493/503. Cells were treated with 75 μM arachidonic acid (AA) for 24 h prior to imaging. Scale bars, 200 μm (zoomed-in, 40 μm). **F** and **G,** Flow cytometry quantification of lipid droplet content in PI3Kβ-null PPB cells and PI3Kβ wild-type PP or PPA cells (**F**), as well as control and CRISPRi-mediated PIK3CB-knockdown BT549 cells (**G**). Cells were treated with 75 μM AA with or without DGAT1 (10 μM) plus DGAT2 (10 μM) inhibitors for 24 h prior to staining and analysis. Data are presented as mean ± s.e.m.; n = 3. **H** and **I,** Heatmap showing z-scored AUC values of RSL3 dose-response curves in PI3Kβ-null PPB cells and PI3Kβ wild-type PP or PPA cells (**H**), as well as control and CRISPRi-mediated PIK3CB-knockdown BT549 cells (**I**). Cells were pretreated with DGAT1 inhibitor (10 μM) plus DGAT2 inhibitor (10 μM) for 24 h prior to 24 h RSL3 treatment. **J,** Model of PI3Kβ-mediated ferroptosis resistance in PTEN-deficient cancers. PI3Kβ promotes lipid metabolic reprogramming, resulting in the accumulation of PUFA-containing neutral lipids and lipid droplet formation. Sequestration of ferroptosis-sensitive PUFAs within lipid droplets limits lipid peroxidation and suppresses ferroptotic cell death.

Quantitative analyses further demonstrated that the lipid species commonly downregulated in PPB relative to PP and PPA cells were predominantly neutral lipids (TAG and CE), with the majority containing PUFAs (**Figure 4C**). Analysis of all identified TAGs and ether-linked TAGs (TAG-O) species revealed that the downregulation in PPB cells was largely confined to PUFA-containing neutral lipids, whereas SFA- or MUFA-containing species were comparatively preserved (**Figure 4D**). Importantly, a similar pattern was observed under ferroptotic stress, where RSL3-treated PI3Kβ-ablated PPB cells exhibited selective depletion of PUFA-containing TAGs and CEs compared with PP and PPA cells (**Figures S6A-S6D**).

Together, these findings indicate that PI3Kβ promotes the accumulation of PUFA-rich neutral lipid stores while limiting the abundance of ferroptosis-sensitive PUFA-containing membrane phospholipids. These results support a model in which PI3Kβ suppresses ferroptosis by channeling PUFAs into lipid droplet-associated storage pools, thereby reducing the availability of lipids susceptible to peroxidation.

### PI3Kβ drives lipid droplet formation to inhibit ferroptosis

Because TAGs and CEs are the principal constituents of lipid droplets, and lipid droplet-mediated PUFA sequestration is a well-established ferroptosis resistance mechanism ^39–41^, we next investigated whether PI3Kβ suppresses ferroptosis by promoting lipid droplet formation and PUFA storage. To test this hypothesis, we stained cells with BODIPY 493/503, a neutral lipid-selective dye that labels lipid droplets. AA treatment robustly induced lipid droplet formation in PP and PPA cells, whereas PI3Kβ-deficient PPB cells exhibited minimal lipid droplet accumulation (**Figure 4E**). Flow cytometry further demonstrated that PI3Kβ depletion or knockdown in both murine and human cancer cells impaired AA-induced lipid droplet formation, whereas PI3Kβ re-expression in PPB cells restored this response (**Figures 4F**, **4G, S6E, and S6F**).

To determine whether PI3Kβ-dependent lipid droplet formation contributes to ferroptosis resistance, we pharmacologically inhibited diacylglycerol O-acyltransferase 1/2 (DGAT1/2), the key enzymes in TAG biosynthesis and lipid droplet formation. DGAT1/2 inhibition abrogated PI3Kβ-driven lipid droplet accumulation (**Figures 4F**, **4G, S6E, and S6F**). Notably, disruption of lipid droplet formation abolished PI3Kβ-mediated ferroptosis resistance in both human and murine PTEN-deficient cancer cells (**Figures 4H**, **4I, S6G, and S6H**), demonstrating that lipid droplet biogenesis is critical for PI3Kβ-dependent ferroptosis protection.

Together with our lipidomic analyses, these findings support a model in which PI3Kβ promotes the sequestration of ferroptosis-sensitive PUFAs into TAG- and CE-rich lipid droplets, thereby limiting the availability of oxidizable membrane phospholipids and preventing lipid peroxidation. Consequently, PI3Kβ-driven lipid droplet formation constitutes a critical mechanism by which PTEN-deficient cancers evade ferroptotic cell death (**Figure 4J**).

### SREBP1 mediates PI3Kβ-driven DNL and ferroptosis resistance

Although PI3K signaling is well established regulator of lipid metabolism ^42^, the isoform-specific transcriptional regulators downstream of PI3Kβ that mediate ferroptosis resistance remain poorly defined. To identify candidate regulators, we analyzed correlations between sensitivity to three ferroptosis inducers, RSL3, ML162, and ML210, and the expression of 19 well-characterized lipid metabolism-associated transcription factors across PTEN-deficient cancer lines in the DepMap dataset. Strikingly, sterol regulatory element-binding transcription factor 1 (*SREBF1*) was the only transcription factor that showed consistent positive association with resistance to all three ferroptosis inducers (**Figures S7A and S7B**). Further analysis revealed that elevated SREBF1 expression was among the strongest predictors of resistance to ML210, ML162, and RSL3 across PTEN-deficient cancer cell lines (**Figure S7C**). Because SREBP1 is a master regulator of DNL, we next investigated whether it is regulated by PI3Kβ signaling. Western blot analyses demonstrated that PI3Kβ activates AKT-mTOR signaling axis and promote SREBP1 activation in both murine and human PTEN-deficient cancer cells (**Figures S7D-S7F**). Collectively, these findings identify SREBP1 as a candidate downstream mediator linking PI3Kβ to ferroptosis resistance.

To investigate whether SREBP1 functions as the downstream effector of PI3Kβ-dependent lipid metabolic reprogramming, we performed RNA-seq analysis in PP and PPB cells expressing empty vector, as well as in PPB cells with ectopic SREBP1 expression. Consistent with our previous findings, PI3Kβ deletion resulted in broad downregulation of lipid metabolic genes (**Figure S8A**). Notably, SREBP1 overexpression in PPB cells reversed these transcriptional defects and induced expression of multiple lipid metabolic genes (**Figure S8B**). Furthermore, GSEA demonstrated that SREBP1 overexpression restored fatty acid metabolism and adipogenesis programs suppressed by PI3Kβ-deficiency (**Figure S8C**).

Given the central role of lipid droplets in PI3Kβ-mediated ferroptosis resistance, we next examined whether SREBP1 regulates lipid droplet formation downstream of PI3Kβ. SREBP1 overexpression completely rescued AA-induced lipid droplet accumulation in PPB cells, whereas treatment with the DGAT1/2 inhibitor abolished this effect (**Figures S8D and S8E**). Similarly, in human BT549 and PC3 cell lines, PI3Kβ knockdown impaired AA-induced lipid droplet formation, which was restored by SREBP1 overexpression but eliminated by inhibition of DGAT1/2 (**Figures S8F and S8G**).

Finally, functional ferroptosis assays showed that SREBP1 restoration largely abrogated the ferroptosis sensitivity induced by PI3Kβ-depletion in both murine and human PTEN-deficient cancer cells (**Figures S8H-S8J**). Together, these findings establish SREBP1 as a critical downstream effector of PI3Kβ signaling that drives DNL, promotes lipid droplet formation, and mediates ferroptosis resistance in PTEN-deficient cancers.

### Therapeutic potential of combined PI3Kβ inhibition and ferroptosis induction

Having established that PI3Kβ suppresses ferroptosis in PTEN-deficient cancers, we next investigated whether this synthetic lethal interaction could be therapeutically exploited for cancer treatment. We first performed Bliss synergy analyses in both murine and human PTEN-deficient cancer cells, comparing the effects of combining ferroptosis inducers (RSL3 or ML210) with either the PI3Kα inhibitor BYL-719 or the PI3Kβ inhibitor AZD6482. Strikingly, AZD6482, but not BYL-719, showed robust synergistic interactions with ferroptosis inducers across all three examined PTEN-deficient cell lines (**Figures S9A-S9E**), further supporting a specific role for PI3Kβ in ferroptosis suppression.

We next evaluated the therapeutic potential of this combination strategy in vivo in PTEN-deficient tumor models. In the PP GEMM, the combination of AZD6482 and imidazole ketone erastin (IKE), the most biologically effective ferroptosis inducer currently available for in vivo studies, synergistically suppressed tumor growth and prolonged survival compared with either monotherapy (**Figures 5A-5C**). Similar antitumor synergy was observed in two human PTEN-deficient xenograft models, PC3 and HCC70, following combined PI3Kβ inhibition and ferroptosis induction (**Figures S10A-S10H**).

**Figure 5.**
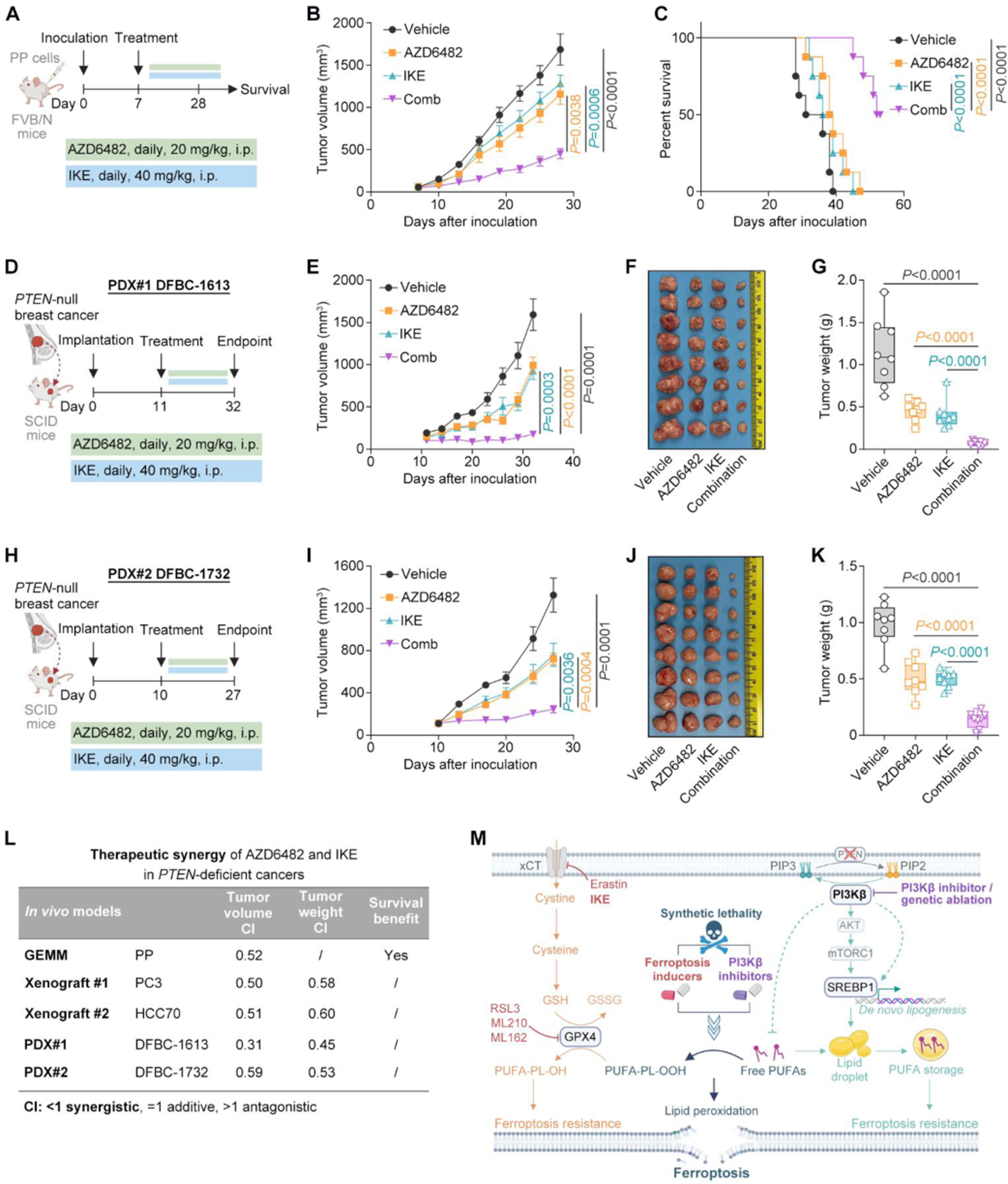
Combined PI3Kβ inhibition and ferroptosis induction synergistically suppress PTEN-deficient tumors. **A-C,** Treatment scheme (**A**), tumor growth curves (**B**), and Kaplan-Meier survival analysis (**C**) of PP GEMMs treated with the indicated drugs. Data are presented as mean ± s.e.m.; two-way ANOVA; n = 8. Survival curves were analyzed using the Kaplan-Meier method with log-rank test. **D-G,** Treatment scheme (**D**), tumor growth curves (**E**), representative tumor images (**F**), and tumor weights (**G**) of the breast cancer PDX model DFBC-1613 treated with the indicated drugs. Data are presented as mean ± s.e.m.; two-way ANOVA for tumor volume; unpaired two-tailed t-test for tumor weight; n = 8. **H-K,** Treatment scheme (**H**), tumor growth curves (**I**), representative tumor images (**J**), and tumor weights (**K**) of the breast cancer PDX model DFBC-1732 treated with the indicated drugs. Data are presented as mean ± s.e.m.; two-way ANOVA for tumor volume; unpaired two-tailed t-test for tumor weight; n = 8. **L,** In vivo synergy analysis using combination index (CI) of AZD6482 and IKE across PTEN-deficient breast cancer GEMMs, breast and prostate cancer xenografts, and breast cancer PDX models. CI was calculated as: CI = (Comb/S2)/(S1/Ctrl), where Ctrl represents the vehicle-treated group, S1 and S2 represent single-agent groups, and Comb represents the combination treatment group. CI < 1 indicates synergy, CI = 1 indicates additivity, and CI > 1 indicates antagonism. **M,** Proposed model for synthetic lethality between PI3Kβ inhibition and ferroptosis induction in PTEN-deficient cancers. PI3Kβ promotes a ferroptosis-resistant state by driving lipid metabolic reprogramming and PUFA sequestration. PI3Kβ inhibition increases PUFA availability, while ferroptosis inducers disrupt the xCT-GSH-GPX4 defense pathway. Together, these effects enhance lipid peroxidation and trigger synergistic cell death.

To further assess the translational potential of this approach, we evaluated the combination in two breast cancer PDX models. Although AZD6482 or IKE monotherapy produced only modest tumor growth inhibition, the combination induced significant and durable tumor suppression in both models (**Figures 5D-5K**). Notably, in the DFBC-1613 model, tumors in the combination group remained near baseline size after one month of treatment, whereas tumors in the control group had almost reached endpoint by the same time point (**Figure 5E**). Quantitative assessment of therapeutic interactions across all five PTEN-deficient models demonstrated strong synergy, with combination index (CI) values ranging from 0.31 to 0.60, and the most pronounced synergy observed in the PDX models (**Figure 5L**).

Collectively, these findings demonstrate that ferroptosis represents a potent therapeutic vulnerability created by PI3Kβ inhibition. The robust efficacy observed across genetically engineered, xenograft, and patient-derived models provides strong preclinical evidence that combined PI3Kβ inhibition and ferroptosis induction may represent an effective therapeutic strategy for PTEN-deficient cancers.

## DISCUSSION

PTEN-deficient tumors represent a substantial proportion of human cancers, yet effective targeted therapeutic strategies for this genetic context remain limited. Previous studies from our group and others established that PTEN-deficient cancers preferentially depend on the PI3Kβ isoform for growth and survival and further demonstrated that PI3Kβ inhibition can activate anti-tumor immunity and synergize with immune checkpoint blockade to achieve significant therapeutic benefit ^5,6,8,12,34^. However, despite the critical role of PI3Kβ in PTEN-deficient tumors, pharmacological PI3Kβ inhibition alone is insufficient to induce complete tumor regression ^8^, highlighting the need to identify effective synthetic lethal vulnerabilities that can be therapeutically exploited. Here, through the integration of large-scale cancer dependency datasets, genetic models, lipidomics, metabolic flux analyses, and functional studies, we identify ferroptosis resistance as a previously unrecognized function of PI3Kβ signaling in PTEN-deficient cancers. Importantly, we demonstrate that PI3Kβ inhibition creates a selective ferroptosis vulnerability that can be therapeutically exploited across multiple preclinical models, including PTEN-deficient PDXs.

Ferroptosis is a distinct form of regulated cell death characterized by iron-dependent lipid peroxidation and is fundamentally different from apoptosis and other forms of cell death ^17^. Extensive efforts have focused on elucidating the molecular mechanisms underlying ferroptosis defense systems, among which the canonical GSH-GPX4 axis ^43,44^ and the FSP1-CoQ_10_ pathway ^45,46^ represent the two major endogenous defense systems. More recently, it has become increasingly clear that distinct oncogenic states can create selective dependencies on specific ferroptosis defense mechanisms, thereby generating targetable therapeutic vulnerabilities. For example, GPX4 dependency has been identified in drug-tolerant persister cells and mesenchymal-like cancer states ^24,25^, whereas FSP1 has emerged as a therapeutic target in lung cancer and metastatic melanoma ^22,47^. Our findings extend this paradigm by identifying PTEN-deficient cancers as a ferroptosis-resistant state that is maintained by PI3Kβ signaling.

Mechanistically, we identify SREBP1-dependent lipid metabolic reprogramming as a key downstream mechanism of PI3Kβ-mediated ferroptosis resistance. Previous studies established that oncogenic PI3K-AKT-mTOR signaling suppresses ferroptosis through activation of SREBP1 and lipogenic programs that limit lipid peroxidation ^30,31,33^. Consistent with this model, we find that PI3Kβ signaling promotes SREBP1 activity and DNL in PTEN-deficient tumors. However, our findings further suggest that PI3Kβ regulates ferroptosis sensitivity through additional mechanisms involving PUFA trafficking and lipid remodeling that extend beyond the AKT-mTOR-dependent transcriptional control of lipogenesis alone.

Through integrated transcriptomic, metabolomic, isotope-tracing, and lipidomic analyses, we show that PI3Kβ promotes lipid droplet formation and the accumulation of PUFA-containing neutral lipids. As a consequence, ferroptosis-sensitive PUFAs are preferentially sequestered into TAG- and CE-rich lipid droplets rather than incorporated into membrane phospholipids that serve as substrates for lipid peroxidation. This redistribution of PUFAs limits the availability of oxidizable membrane lipids and thereby suppresses ferroptotic cell death. Thus, PI3Kβ establishes a lipid storage state that protects PTEN-deficient tumors from ferroptotic cell death.

More broadly, our work supports an emerging model in which oncogenic signaling pathways suppress ferroptosis through metabolic rewiring. In PTEN-deficient context, PI3Kβ signaling appears to regulate not only lipogenic gene expression through the AKT-mTOR-SREBP1 axis but also the partitioning of PUFAs into neutral lipid storage pools through mechanisms that may be at least partially independent of AKT-mTOR signaling. One intriguing possibility is that PI3Kβ influences AA partitioning through phosphoinositide metabolism. Phosphoinositides serve as important reservoirs of AA and participate in lipid remodeling pathways that may influence ferroptosis sensitivity. Whether PI3Kβ directly regulates these processes remains an important question for future investigation. Thus, our findings propose lipid metabolism as a central determinant of ferroptosis competence and reveal multiple levels at which oncogenic signaling can modulate ferroptotic responses.

These findings have important therapeutic implications. We show that inhibition of PI3Kβ disrupts PUFA sequestration and renders PTEN-deficient tumors highly dependent on ferroptosis defense pathways for survival. As a result, co-targeting PI3Kβ and ferroptosis defense mechanisms produces robust synergistic anti-tumor activity across GEMMs, human xenografts, and PDXs. Notably, the selective synergy observed with PI3Kβ inhibition, but not PI3Kα inhibition, further highlights the unique role of PI3Kβ in governing ferroptosis resistance in PTEN-deficient cancers. More generally, these results suggest that inhibition of oncogenic signaling pathways can create therapeutically exploitable ferroptosis vulnerabilities by disrupting metabolic programs that normally protect tumor cells from lipid peroxidation. In PTEN-deficient cancers, ferroptosis induction may therefore provide an effective strategy to overcome the limited efficacy of PI3Kβ-targeted monotherapy.

In summary, our study identifies PI3Kβ as a key determinant of ferroptosis resistance in PTEN-deficient cancers and establishes a mechanistic axis linking PTEN loss, PI3Kβ-driven lipid metabolic reprogramming, and ferroptosis suppression. Our findings also suggest that PI3Kβ may influence PUFA metabolism and lipid storage through multiple downstream pathways, offering additional insight into how oncogenic PI3K signaling shapes ferroptosis sensitivity. These findings provide a mechanistic rationale for combining PI3Kβ inhibitors with ferroptosis-inducing therapies as a precision therapeutic strategy for PTEN-deficient tumors (**Figure 5M**).

## METHODS

### Cell culture

Mouse primary mammary tumor cell lines (PP, PPA, and PPB) derived from genetically engineered mouse models (GEMMs) were isolated from orthotopic mammary tumor models established in our previous study and cultured in vitro according to previously described protocols ^8,34^. HEK293T (CRL-3216), human breast cancer cell lines HCC70 (CRL-2315) and BT549 (HTB-122), and the human prostate cancer cell line PC3 (CRL-1435) were obtained from the American Type Culture Collection. HEK293T cells were maintained in Dulbecco’s Modified Eagle Medium (DMEM) supplemented with 10% fetal bovine serum (FBS), HCC70, BT549, and PC3 cells were cultured in RPMI-1640 medium supplemented with 10% FBS. All cell cultures were supplemented with 100 μg ml^-1^ penicillin-streptomycin (15140122, Gibco) and maintained at 37 °C in a humidified incubator with 5% CO_2_. All human cell lines were authenticated by short tandem repeat profiling. All cells were routinely tested and confirmed to be free of mycoplasma contamination prior to experimentation.

### Animal studies

All animal experiments were approved by the Institutional Animal Care and Use Committee (IACUC) of Dana-Farber Cancer Institute (protocol 03-111) and conducted in accordance with institutional guidelines. All experimental endpoints complied with approved ethical regulations. Tumor volume was calculated using the formula: 0.52 × length × width^2^.

#### GEMM-derived tumor model

Mouse mammary tumor cells (PP) were orthotopically implanted (5 × 10^5^ cells) into the mammary fat pads of 6-week-old female FVB/N mice (001800, The Jackson Laboratory). When tumors reached approximately 50-100 mm^3^, mice were randomly assigned to treatment groups. Tumor growth was monitored by caliper measurements, and overall survival was recorded.

#### Cell-derived xenografts

Human breast cancer HCC70 cells (3 × 10^6^ cells) were orthotopically implanted into the mammary fat pads of 6-week-old female SCID mice (ICRSC-F, Taconic). Human prostate cancer PC3 cells (1 × 10^6^ cells) were implanted subcutaneously into 6-week-old male SCID mice (ICRSC-M, Taconic). When tumors reached approximately 50-100 mm^3^, mice were randomized for treatment and tumor growth was monitored. For inducible GPX4 knockdown experiments, mice in the treatment group were administered doxycycline via a doxycycline-containing chow diet (2000 ppm), whereas control mice received standard chow without doxycycline.

#### Patient-derived xenografts

Breast cancer patient tumor samples were obtained as previously described^8^, with informed consent from patients. Tumor fragments were implanted into 6-week-old female SCID mice (ICRSC-F, Taconic). Drug treatment was initiated once tumors were established and exhibited stable growth (approximately 2 weeks after implantation), and tumor volume was monitored thereafter.

#### Drug treatment

Mice were treated with the following regimens: AZD6482: 20 mg kg^-1^, intraperitoneally, once daily; IKE: 40 mg kg^-1^, intraperitoneally, once daily; and Liproxstatin-1 (Lip-1): 10 mg kg^-1^, intraperitoneally, once daily.

#### In vivo combination index analysis

The combination index (CI) for in vivo studies was calculated as: CI = (Comb/S2)/(S1/Ctrl), where Ctrl represents the vehicle-treated group, S1 and S2 represent single-agent treatment groups, and Comb represents the combination treatment group. CI < 1 indicates synergy, CI = 1 indicates an additive effect, and CI > 1 indicates antagonism.

### Cloning and lentivirus generation

Lentiviruses were produced in HEK293T cells using a second-generation packaging system consisting of psPAX2 (#12260, Addgene) and pMD2.G (#12259, Addgene). For doxycycline-inducible GPX4 knockdown, the Tet-pLKO-puro vector was originally developed by the Novartis Institutes for Biomedical Research and later deposited to Addgene (#21915), and then shRNA sequences targeting GPX4 were cloned into the Tet-pLKO-puro vector. For CRISPRi-mediated gene silencing, cells were first transduced with lentivirus encoding pLX_311-KRAB-dCas9 (#96918, Addgene) and selected with blasticidin (6 μg ml^-1^; R21001, Thermo Fisher Scientific) for approximately 2 weeks to establish stable KRAB-dCas9-expressing cell lines. Subsequently, gRNA sequences targeting genes of interest were cloned into the pXPR_050 vector (#96925, Addgene) and introduced into KRAB-dCas9-expressing cells via lentiviral transduction. Cells were then selected with puromycin (1 μg ml^-1^) for 1 week to generate stable knockdown cell lines. For lentivirus-based PI3Kβ knockdown and overexpression, constructs from our previously established system were used^8^. SREBP1 overexpression plasmids pLV-puro-N-3HA-SREBP1-C-Flag (#134286) and pLX_TRC317-SREBP1 (#143645) were obtained from Addgene. The human GPX4 shRNA sequence (5’ to 3’) is: GTGGATGAAGATCCAACCCAA. CRISPRi gRNA sequences were selected from the Dolcetto CRISPRi library targeting the indicated genes ^48^.

### Reagents

The following compounds were used in this study: RSL3 (S8155, Selleck Chemicals), ML210 (SML0521, Sigma-Aldrich), ML162 (S4452, Selleck Chemicals), Erastin (S7242, Selleck Chemicals), Bortezomib (S1013, Selleck Chemicals), Paclitaxel (S1150, Selleck Chemicals), Elesclomol (S1052, Selleck Chemicals), Zinc pyrithione (S4075, Selleck Chemicals), Polyphyllin VI (S9302, Selleck Chemicals), Nigericin sodium salt (S6653, Selleck Chemicals), Doxycycline (HY-N0565, MedChemExpress), AZD6482 (HY-10344, MedChemExpress), BYL-719 (HY-15244, MedChemExpress), Liproxstatin-1 (S7699, Selleck Chemicals), Belnacasan (VX-765; S2228, Selleck Chemicals), Ammonium tetrathiomolybdate (TTM; 323446, Sigma-Aldrich), Z-VAD-FMK (S7023, Selleck Chemicals), Deferasirox (S1712, Selleck Chemicals), Ferrostatin-1 (S7243, Selleck Chemicals), DGAT1 inhibitor 4a (T6365, TargetMol), PF-06424439 methanesulfonate (T12425, TargetMol), and Imidazole ketone erastin (IKE; T5523, TargetMol). The following metabolites were used: Sodium palmitate (P9767, Sigma-Aldrich), Arachidonic acid (10931, Sigma-Aldrich), and D-glucose (6-^13^C, 99%; CLM-481, Cambridge Isotope Laboratories).

### Cell viability analysis

Cell viability was assessed using the CellTiter-Glo 2.0 Cell Viability Assay (G9242, Promega) according to the manufacturer’s instructions. For cell death inducer screening, cells were treated with compounds at a range of concentrations for 24 h, followed by measurement of cell viability. Dose-response curves were generated and the area under the curve (AUC) was calculated using GraphPad Prism 9. For Bliss synergy analysis, cells were treated with a matrix of two drugs for 48 h, after which cell viability was measured. Bliss synergy scores were calculated using SynergyFinder 3.0 ^49^.

### Western blot

Cells were lysed in RIPA buffer (89901, Thermo Fisher Scientific) supplemented with protease inhibitors. Protein samples were denatured at 99 °C for 10 min prior to loading. Equal amounts of protein were separated by SDS-PAGE using gels ranging from 6% to 15%, depending on the molecular weight of the target proteins, and subsequently transferred onto PVDF or nitrocellulose membranes. Membranes were blocked with 5% BSA at room temperature for 1 h and then incubated with primary antibodies overnight at 4 °C. After washing, membranes were incubated with fluorescence-conjugated secondary antibodies for 2 h at room temperature. Protein bands were visualized using an Odyssey imaging system (LI-COR Biosciences). The following primary antibodies were used: anti-GPX4 (1:1000; 67763-1-Ig, Proteintech), anti-p110β (1:1000; 3011S, Cell Signaling Technology), anti-phospho-AKT (Ser473) (1:1000; 9271S, Cell Signaling Technology), anti-AKT (1:1000; 9272S, Cell Signaling Technology), anti-phospho-mTOR (Ser2448) (1:1000; 2971S, Cell Signaling Technology), anti-mTOR (1:1000; 2972S, Cell Signaling Technology), anti-phospho-p70 S6K (Thr389) (1:1000; 9234S, Cell Signaling Technology), anti-p70 S6K (1:1000; 9202S, Cell Signaling Technology), anti-SREBP1 (1:1000; ab28481, Abcam), and anti-β-actin (1:3000; A2228, Sigma-Aldrich). Secondary antibodies included anti-mouse IgG (1:2000; RL610-145-002, Rockland Immunochemicals) and anti-rabbit IgG (1:2000; 35568, Thermo Fisher Scientific).

### Flow cytometry

#### Lipid peroxidization detection

Cells were treated with 500 nM RSL3 or 1 μM ML210 at 37 °C for 3 h. Following treatment, cells were harvested by trypsinization, neutralized, and pelleted by centrifugation. Cell pellets were resuspended in PBS supplemented with 1% FBS containing 5 μM BODIPY^TM^ 581/591 C11 (D3861, Thermo Fisher Scientific) and incubated at 37 °C for 30 min. After staining, cells were washed twice with PBS containing 1% FBS and resuspended in the same buffer for flow cytometric analysis. Lipid peroxidation was assessed by measuring fluorescence intensity in the FITC channel (oxidized lipid species) and PE channel (reduced/non-oxidized lipid species).

#### Lipid droplet quantification

Cells were seeded at a density of approximately 2 × 10^5^ cells per well in 6-well plates. After 12 h of attachment, cells were treated with 75 μM arachidonic acid (AA) in the presence or absence of DGAT1 and DGAT2 inhibitors (10 μM each) for 24 h. Cells were then harvested by trypsinization, neutralized, and pelleted by centrifugation. Cell pellets were resuspended in PBS containing 1% FBS and stained with 5 μM BODIPY^TM^ 493/503 (D3922, Thermo Fisher Scientific) at 37 °C for 30 min. After washing twice with PBS containing 1% FBS, cells were resuspended in the same buffer and subjected to flow cytometry. Fluorescence intensity in the FITC channel was used as a measure of neutral lipid content, representing intracellular lipid droplets. Samples were acquired on an LSR Fortessa HTS flow cytometer (BD Biosciences), with data collected using BD FACSDiva software and analyzed using FlowJo.

### Morphological analysis

#### Bright-field imaging

Cells were seeded in 6-well plates and subjected to the indicated treatments. Bright-field images were acquired using a Cytation imaging system (BioTek).

#### Lipid droplet imaging

For lipid droplet fluorescence imaging, cells were stained with 5 μM BODIPY^TM^ 493/503 (D3922, Thermo Fisher Scientific) diluted in PBS at 37 °C for 30 min. After staining, cells were washed twice with PBS and maintained in PBS during imaging. Fluorescence images were acquired using a Cytation imaging system to visualize lipid droplets (green channel). Corresponding phase-contrast images were captured to define cell morphology and localization.

### Fatty acid oxidation assay

Fatty acid oxidation was assessed using the Long Chain Fatty Acid Oxidation Stress Test Kit (103672-100, Agilent Technologies) according to the manufacturer’s instructions on a Seahorse XFe96 Analyzer (Agilent Technologies). Palmitate was used as the substrate and added to the assay medium at a final concentration of 200 μM. Etomoxir (ETO) was injected at a final concentration of 4 μM. Oxygen consumption rate (OCR) was measured following sequential injections according to the mitochondrial stress test protocol.

### Metabolomics and lipidomics

Untargeted metabolomics and lipidomics analyses were performed at the Dana-Farber Cancer Institute Metabolomics Core. For metabolomics, cells were cultured in 6-well plates to approximately 70% confluence and subjected to the indicated treatments. Prior to extraction, cells were washed three times with 75 mM ammonium carbonate buffer (pH 7.4). Metabolites were extracted using a pre-chilled solvent containing acetonitrile:methanol:water (40:40:20, v/v/v). Specifically, 400 μl of extraction solvent was added to each well and incubated at −20 °C for 10 min, followed by collection of the extract. This step was repeated once, and a total of 800 μl extract was pooled per sample. Extracts were centrifuged at 14,000 rpm for 10 min at 4 °C, and the supernatants were collected and stored at −80 °C until analysis. For lipidomics, lipids were extracted using 90% isopropanol (IPA) with SPLASH Lipidomix (330731, Avanti Polar Lipids) added as internal standards. Metabolite identification was performed using two complementary approaches: (i) data-dependent acquisition, in which MS/MS spectra were matched against publicly available spectral libraries based on cosine similarity scores and fragment matching; and (ii) a standard library approach, where metabolites were identified by comparison to a reference library of approximately 1,300 authentic compounds with matched retention times and MS/MS spectra. For lipidomics, lipid species were first annotated based on MS/MS spectra, followed by confirmation using MS1 features with high-confidence signal matching. Each experimental group included four biological replicates, along with one parallel sample used for cell counting and normalization. Differential metabolite analysis and pathway enrichment analysis were performed using the MetaboAnalyst online platform ^50^.

### Stable isotope tracing

Stable isotope tracing was performed to assess de novo lipogenesis. Cells were cultured in 6-well plates and incubated with D-glucose (U-^13^C_6_, 99%; CLM-1396, Cambridge Isotope Laboratories) at a final concentration of 4.5 g l^-1^ for either 1 h (short-term labeling) or 24 h (long-term labeling). Following labeling, cells were washed three times with 75 mM ammonium carbonate buffer (pH 7.4) and metabolites were extracted using pre-chilled acetonitrile:methanol:water (40:40:20, v/v/v) at −20 °C. Isotopic tracing detection and mass distribution vector (MDV) analysis were performed by General Metabolics. Analytes were separated using reverse-phase chromatography and analyzed in negative ionization mode to facilitate fatty acid detection. To define the detectable fatty acid pool, a pooled reference sample generated from unlabeled samples was analyzed to annotate fatty acids based on MS1 features and retention time matching against a column-specific reference library. For each annotated fatty acid, all theoretically possible isotopologues were computed, and the retention time of the unlabeled species was used to assign labeled counterparts. Experimental samples were acquired using an MS1 scanning method, and both labeled and unlabeled isotopologues were identified based on MS1 and retention time matching. The incorporation of ^13^C into fatty acids was quantified, and differences in isotopic labeling patterns were compared across conditions and time points. Each experimental group included four biological replicates, along with one parallel sample used for cell counting and normalization.

### RNA-seq and data analysis

RNA-seq experiments in this study were performed by Novogene. Total RNA was subjected to quality control, and poly(A)+ mRNA was enriched using oligo(dT) magnetic beads, followed by fragmentation and cDNA library construction. Libraries were prepared through standard procedures including end repair, A-tailing, adaptor ligation, PCR amplification, and purification, and sequenced on an Illumina platform. Raw reads were filtered to remove adaptor sequences, reads containing >10% ambiguous bases, and low-quality reads. Clean reads were aligned to the mouse reference genome (mm39) using HISAT2 (v2.2.1) with default parameters. Gene expression levels were quantified using featureCounts (v2.0.6) and normalized as fragments per kilobase of transcript per million mapped reads (FPKM). Differential gene expression analysis was performed using DESeq2 (v1.42.0). Genes with fold change ≥ 2 and p-value ≤ 0.05 were considered significantly differentially expressed. The RNA-seq data generated in this study have been deposited in the Gene Expression Omnibus (GEO) under accession number GSE329097. Gene set enrichment analysis (GSEA) was performed using GSEA software (v4.3.2). For PP, PPA, and PPB tumor RNA-seq datasets, as well as RNA-seq data from PP GEMM tumors treated with vehicle or AZD6482 and PDX DFBC-1807 tumors treated with vehicle or AZD6482, gene expression data were obtained from our previous study and reanalyzed by GSEA.

### CRISPR dependency map analysis

All DepMap correlation analyses were performed using a custom analysis pipeline in the portal based on Pearson correlation. Gene effect data were obtained from DepMap Public 24Q4 and 25Q2 (Chronos scores). Gene expression data were derived from the Expression Public 24Q4, 25Q2 and 26Q1 datasets. ssGSEA scores were obtained from the ssGSEA Public 24Q2 dataset.

### Drug sensitivity analysis

Correlation analyses between gene expression (*PTEN*, *PIK3CB*, and *SREBF1*) and drug sensitivity were performed using the DepMap portal and the Cancer Therapeutics Response Portal online platform. Gene expression data were obtained from Expression Public 24Q4 and 26Q1, drug sensitivity data were derived from the Cancer Target Discovery and Development (CTD2) dataset and the PRISM Repurposing Public 24Q2 dataset.

### Quantification and statistical analysis

Statistical analyses between two groups were performed using two-sided t-tests. Differences in cell viability curves or tumor growth curves were assessed using two-way ANOVA. Survival analyses were performed using the log-rank test. All analyses were conducted using GraphPad Prism 9. DepMap correlation analyses were performed using Pearson correlation. Data are presented as mean ± s.d. or mean ± s.e.m., as indicated in the figure legends. P value ≤ 0.05 was considered statistically significant, and at least three independently experiments with similar results were reported.

### Data availability

The RNA-seq data generated in this study have been deposited in the Gene Expression Omnibus (GEO) under accession number GSE329097. The RNA-seq data of PP, PPA, and PPB tumor, PP GEMM tumors treated with vehicle or AZD6482, and PDX DFBC-1807 tumors treated with vehicle or AZD6482 were obtained from our previously published study (GEO accession: GSE225064)^8^. Pan-cancer cell line gene expression and dependency data were obtained from the Cancer Dependency Map. Gene effect data were derived from DepMap Public 24Q4 and 25Q2 (Chronos scores), and gene expression data were obtained from the Expression Public 24Q4, 25Q2 and 26Q1 datasets. Drug sensitivity data were obtained from the PRISM Repurposing Public 24Q2 dataset and the Cancer Therapeutics Response Portal (CTRP).

## DECLARATION OF INTERESTS

L.C.C. is a founder and member of the Board of Directors of Agios Pharmaceuticals, a founder of Petra Pharmaceuticals, and a co-founder and shareholder of Faeth Therapeutics. J.J.Z. is a co-founder and board director of Crimson Biotech Inc. and Geode Therapeutics Inc. All other authors declare no competing interests.

## AUTHOR CONTRIBUTIONS

Y.N. and J.J.Z. conceived and designed the study. Y.N. performed the majority of the experiments. T.J., J.N., F.C., Y.W., and W.W. assisted with animal experiments. L.C.C. contributed to discussions and provided constructive insights on PI3K biology. J.J.Z. supervised the overall research and experimental execution. Y.N. drafted the manuscript, and J.J.Z. reviewed and finalized the manuscript.

## ACKNOWLEDGMENTS

This work was supported by the Breast Cancer Research Foundation BCRF-24-179 (J.J.Z.), and the National Health Institute (NIH) P50 CA168504 (J.J.Z), P01 CA269021 (J.J.Z.) and CA210057 (J.J.Z). The graphical abstract was created by BioRender.com.

---

**Figure S1.**
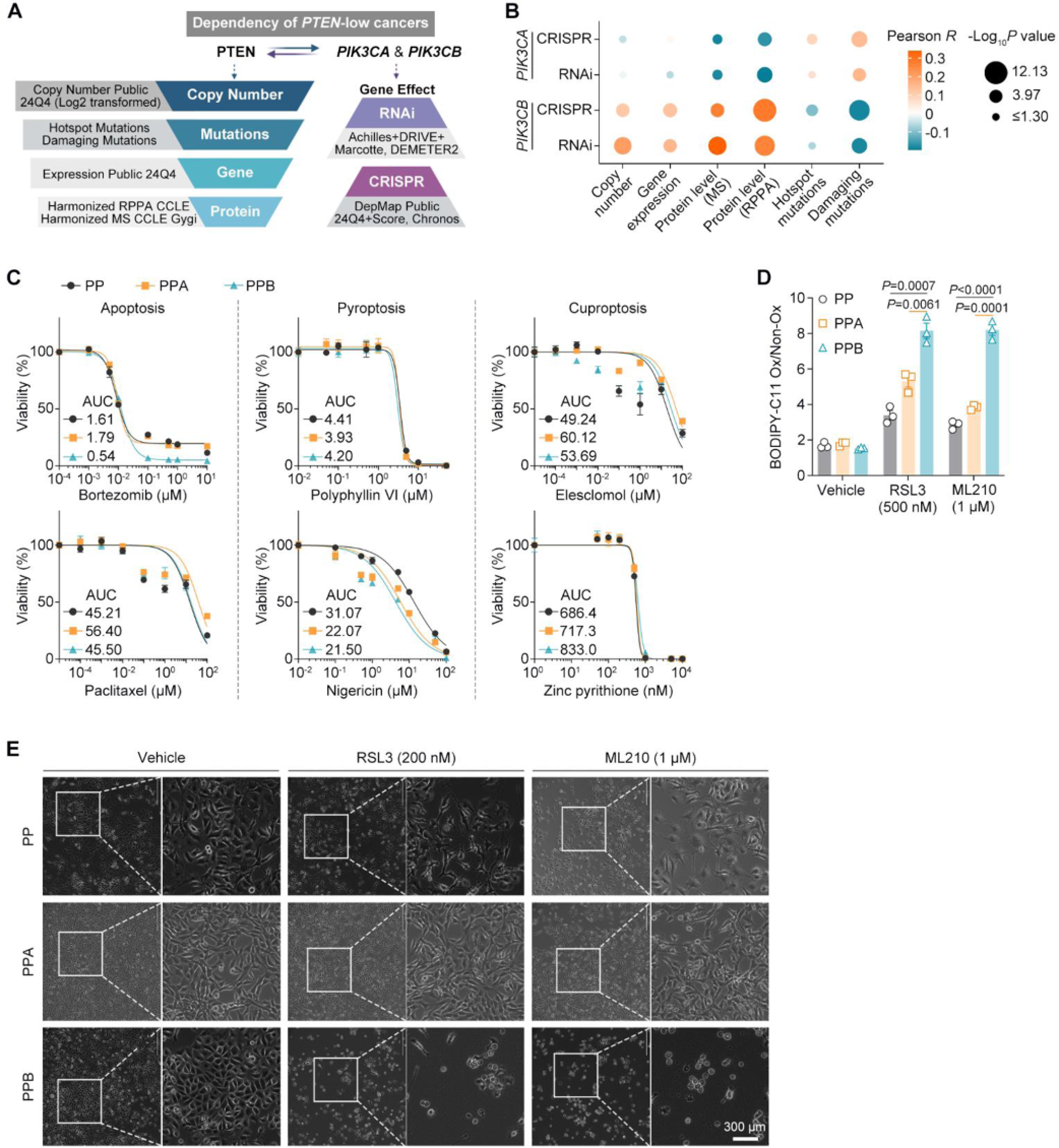
PI3Kβ deletion creates a selective ferroptosis vulnerability in PTEN-deficient cancers. **A,** Schematic illustrating the DepMap analysis strategy to assess correlations between PTEN status (CNV, mutations, mRNA expression, and protein levels) and dependency on PIK3CA and PIK3CB. **B,** Bubble plot showing Pearson correlations between PTEN status and gene effect scores of PIK3CA and PIK3CB. **C,** Dose-response curves of GEMM-derived primary breast cancer cells treated with the indicated cell death inducers. Data are presented as mean ± s.d.; n = 4. **D,** Lipid peroxidation levels in PP, PPA, and PPB cells following RSL3 or ML210 treatment. Data are presented as mean ± s.e.m.; unpaired two-tailed t-test; n = 3. **E,** Bright-field images showing morphological changes in PP, PPA, and PPB cells after 24 h treatment with RSL3 or ML210.

**Figure S2.**
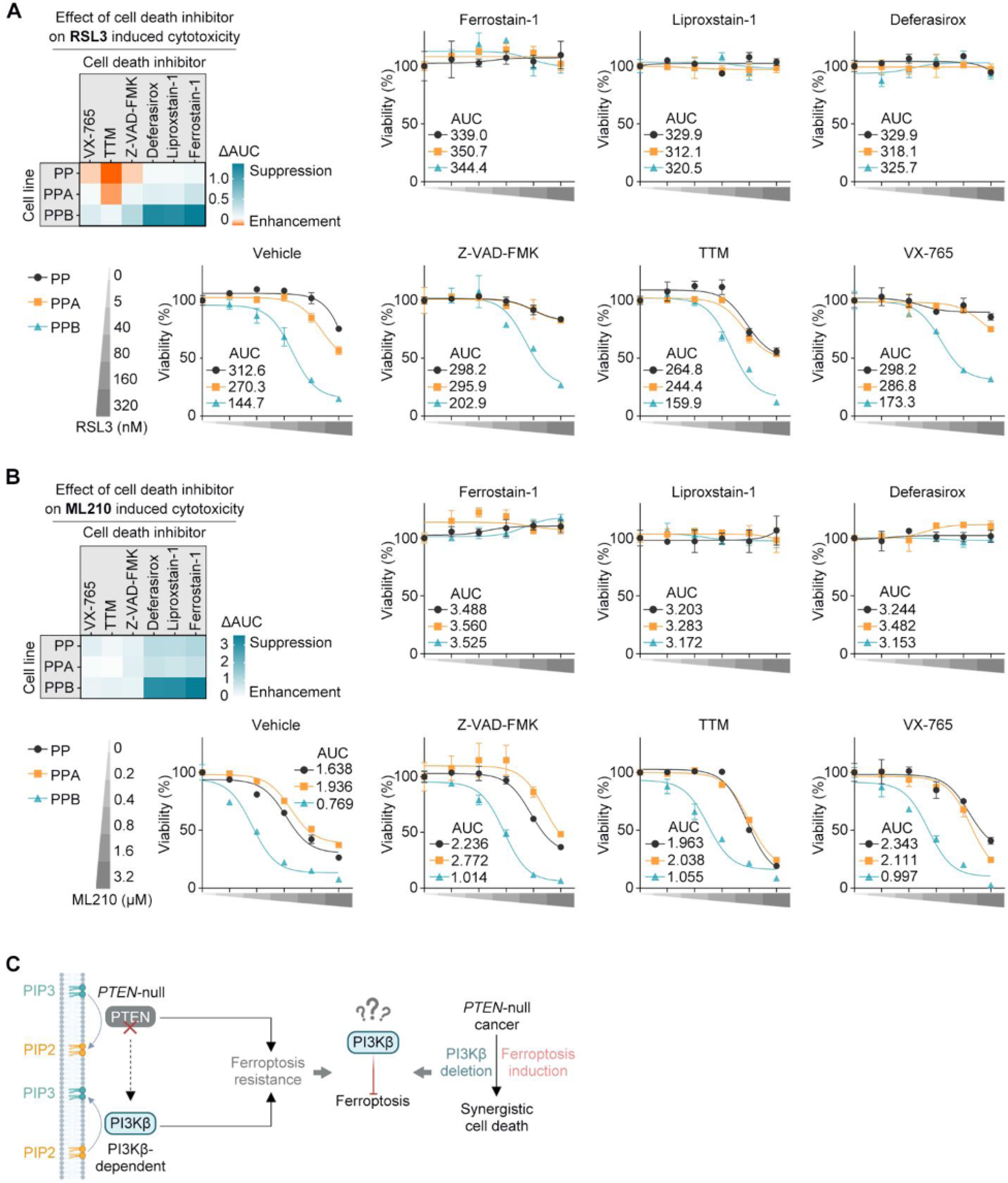
PI3Kβ loss selectively sensitizes PTEN-deficient cancer cells to ferroptosis. **A** and **B,** Heatmaps of AUC changes and dose-response curves following treatment with RSL3 (24 h) or ML210 (48 h). Cells were pretreated with the indicated cell death inhibitors for 24 h prior to RSL3 or ML210 treatment. Data are presented as mean ± s.d.; n = 3. **C,** Schematic model showing that PTEN-deficient cancers are resistant to ferroptosis, whereas PI3Kβ inhibition synergizes with ferroptosis to induce cell death in PTEN-deficient cancers, supporting the hypothesis that PI3Kβ inhibits ferroptosis.

**Figure S3.**
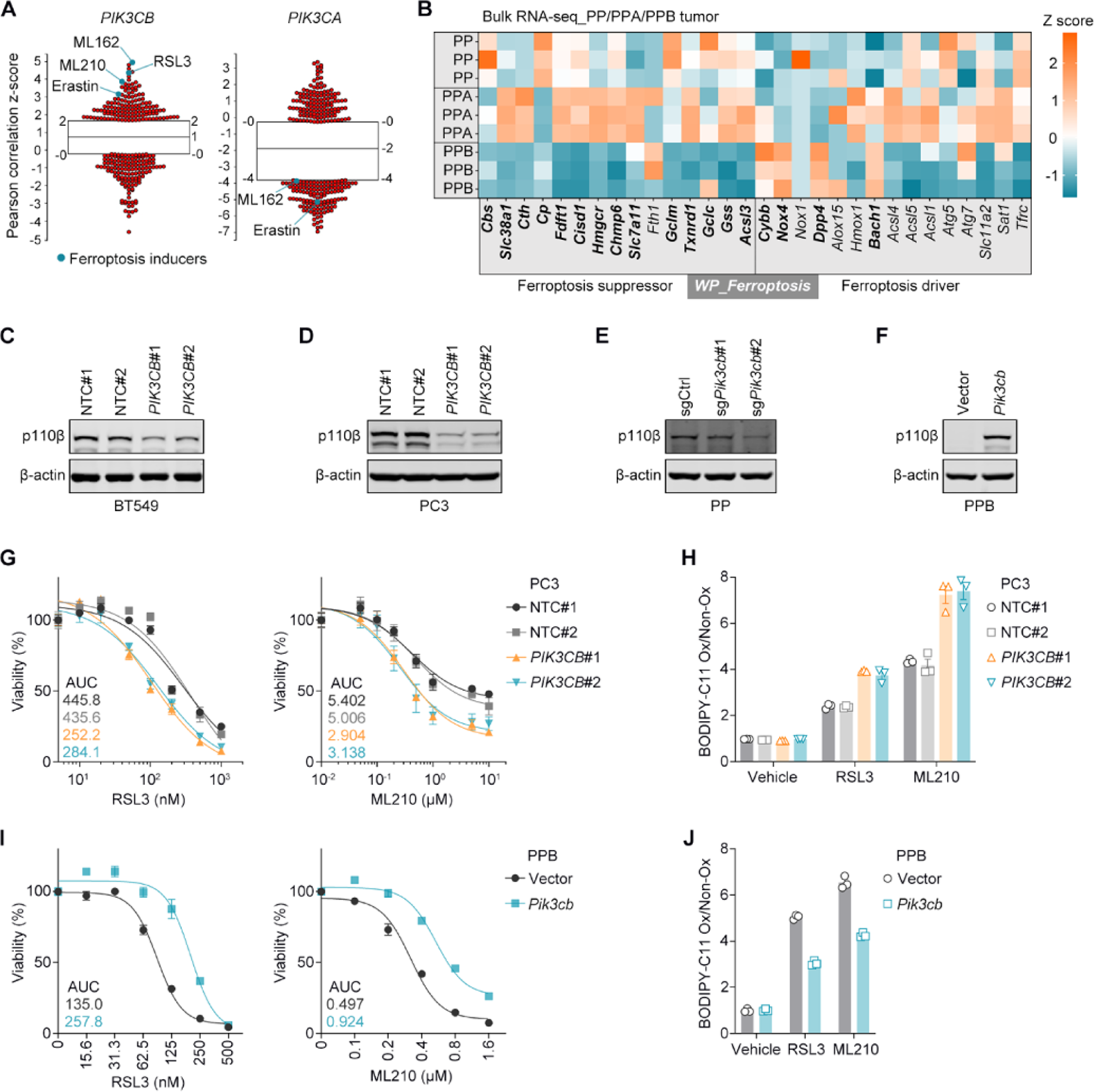
Genetic and pharmacologic evidence that PI3Kβ suppresses ferroptosis. **A,** Correlation analysis of PIK3CB and PIK3CA expression with compound drug sensitivity across pan-cancer cell lines from the CTRP database. Ferroptosis inducers are highlighted in green. Plotted values represent z-scored Pearson correlation coefficients. **B,** Heatmap showing expression of ferroptosis-sensitive and ferroptosis-resistant genes in RNA-seq data from PP, PPA, and PPB tumors. Gene sets were derived from the WP_Ferroptosis pathway. **C-F,** Western blot analysis of p110β protein levels in the indicated cells. **G** and **I,** Dose-response curves of RSL3 (24 h) and ML210 (48 h) in the indicated cells. Data are presented as mean ± s.d.; n = 3. **H** and **J,** Lipid peroxidation levels in the indicated cells following RSL3 or ML210 treatment. Data are presented as mean ± s.e.m.; unpaired two-tailed t-test; n = 3.

**Figure S4.**
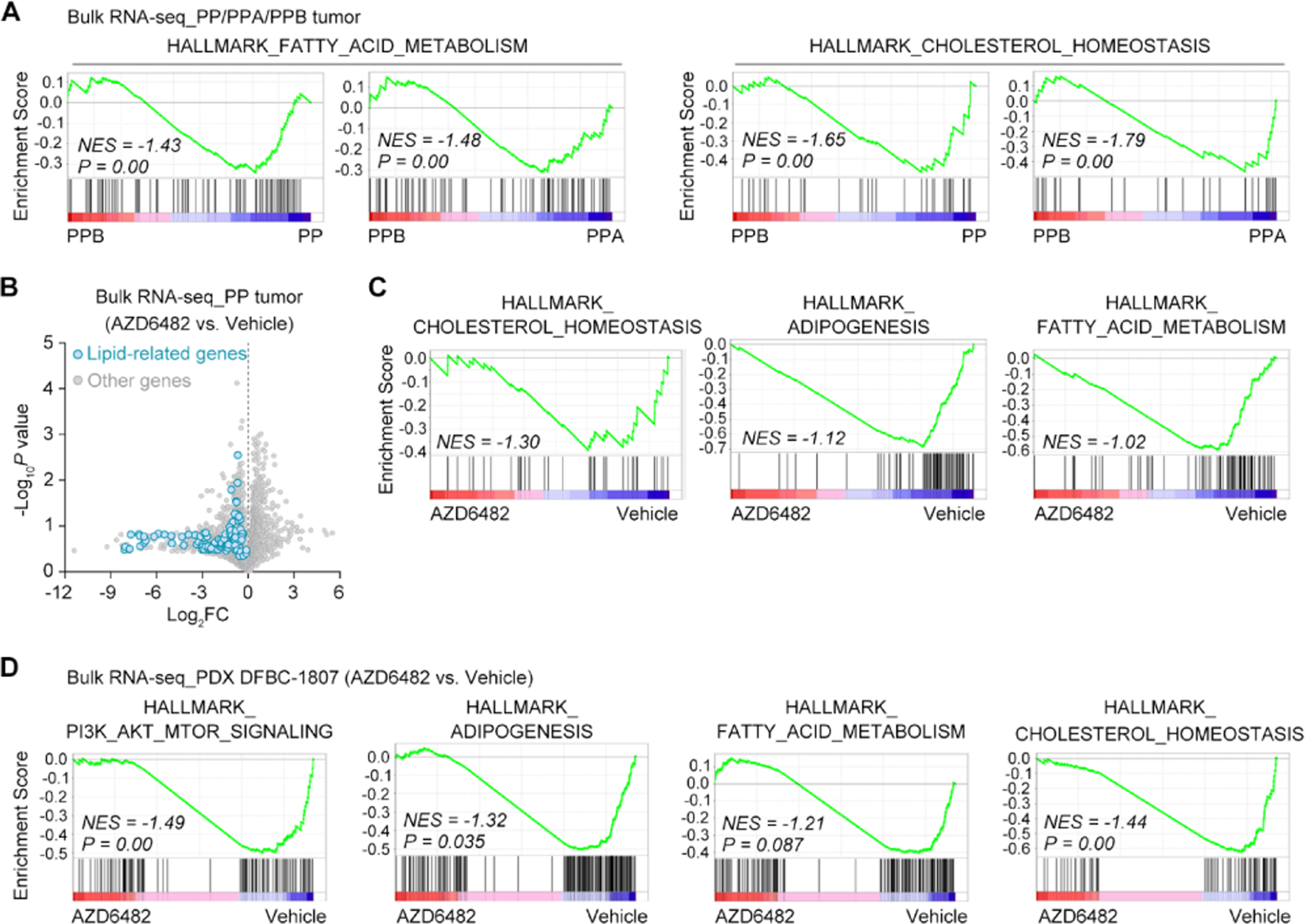
PI3Kβ reprograms lipid metabolism in PTEN-deficient cancers. **A,** GSEA results comparing RNA-seq data from PPB tumors versus PP or PPA tumors. **B,** Volcano plot of differentially expressed genes from RNA-seq data of PP tumors treated with AZD6482 or vehicle. Lipid metabolism-related genes are highlighted in green. **C** and **D,** GSEA of RNA-seq data from PP tumors (**C**) and breast cancer PDX DFBC-1807 tumors (**D**) treated with AZD6482 or vehicle.

**Figure S5.**
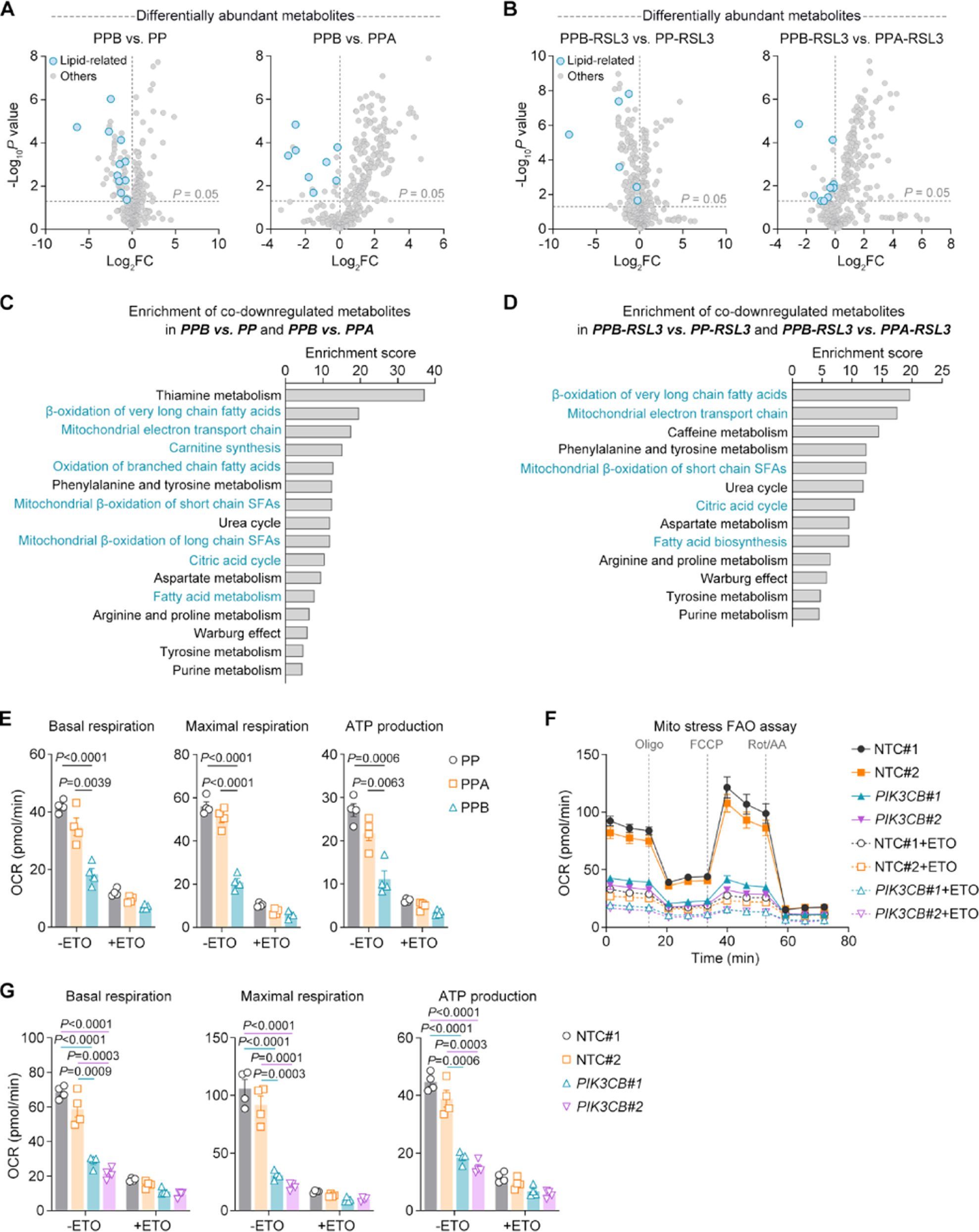
PI3Kβ rewires lipid metabolism in PTEN-deficient cancers. **A** and **B,** Volcano plots showing differentially abundant metabolites from untargeted metabolomics of PP, PPA, and PPB cells with or without RSL3 treatment. Lipid metabolism-related metabolites are highlighted in green. **C** and **D,** Pathway enrichment analysis of commonly downregulated metabolites in PPB cells compared to PP or PPA cells, with or without RSL3 treatment. **E-G,** Oxygen consumption rate (OCR)-based analysis of fatty acid oxidation in the indicated cells using Seahorse assays. Data are presented as mean ± s.e.m.; unpaired two-tailed t-test; n = 4.

**Figure S6.**
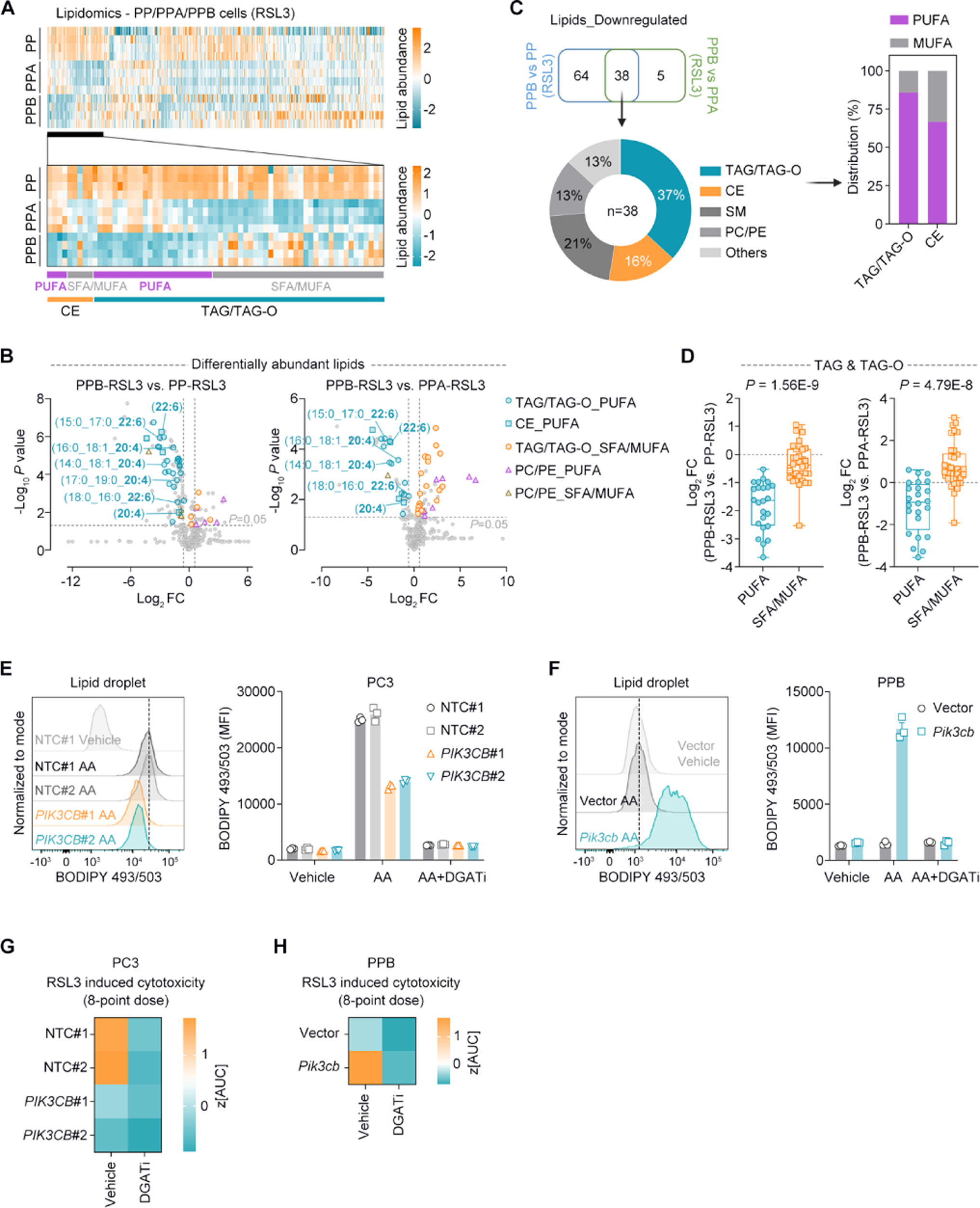
PI3Kβ suppresses ferroptosis by promoting lipid droplet formation and PUFA sequestration. **A,** Heatmap of lipid abundance from lipidomics profiling of RSL3-treated PP, PPA, and PPB cells. **B,** Volcano plots of differentially enriched lipids in RSL3-treated PPB versus PP or PPA cells. PUFA-containing triglycerides (TAG) and cholesteryl esters (CE) are significantly reduced in PPB cells. **C,** Venn diagram showing 38 commonly downregulated lipids in RSL3-treated PPB versus PP and PPB versus PPA. A pie chart indicates that 53% of these lipids are TAG/TAG-O and CE, with bar plots showing the distribution of PUFA- and MUFA-containing species. **D,** Fold-change analysis of PUFA- and SFA/MUFA-containing TAG/TAG-O species. **E** and **F,** Flow cytometry quantification of lipid droplets. Cells were treated with 75 μM arachidonic acid (AA) with or without DGAT1 and DGAT2 inhibitors for 24 h. Data are presented as mean ± s.e.m.; n = 3. **G** and **H,** Heatmaps showing z-scored AUC values of RSL3 dose-response curves in the indicated cells following DGAT1/2 inhibitor pretreatment.

**Figure S7.**
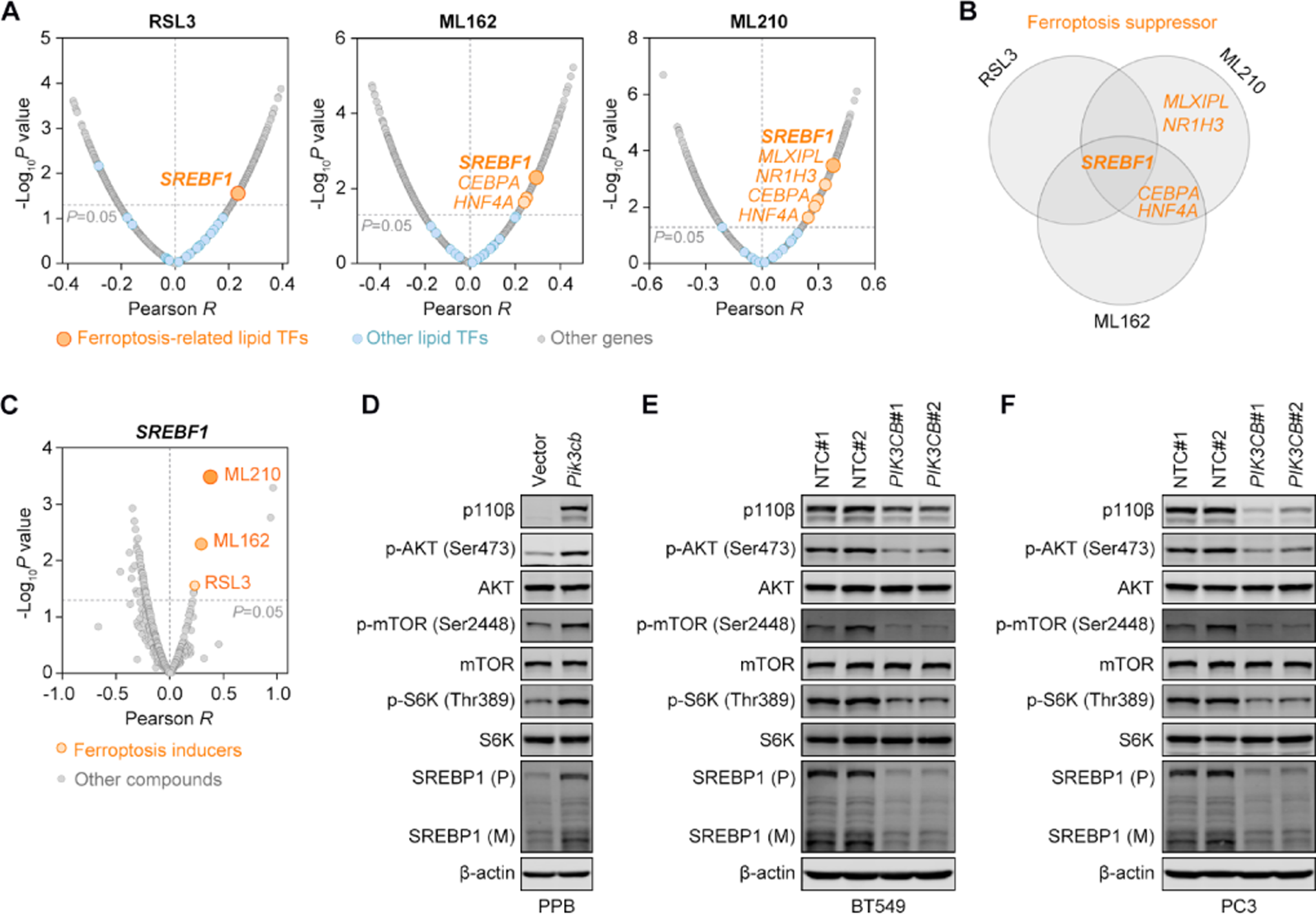
SREBP1 is a candidate downstream effector of PI3Kβ signaling to ferroptosis resistance in PTEN-deficient cancers. **A,** Pearson correlation analysis between gene expression and drug sensitivity AUC values for the ferroptosis inducers RSL3, ML162, and ML210 in PTEN-deficient cancer cell lines. Lipid metabolism-related transcription factors significantly positively associated with ferroptosis resistance are highlighted in yellow, while other lipid metabolism-associated transcription factors are labeled in green. **B,** Venn diagram showing the overlap of lipid metabolism-related transcription factors associated with resistance to all three ferroptosis inducers. **C,** Pearson correlation analysis between SREBF1 expression and compound sensitivity across PTEN-deficient cancer cell lines. Ferroptosis inducers are highlighted in yellow. **D-F,** Western blot analysis of PI3K-AKT signaling and SREBP1 protein levels in the indicated cells.

**Figure S8.**
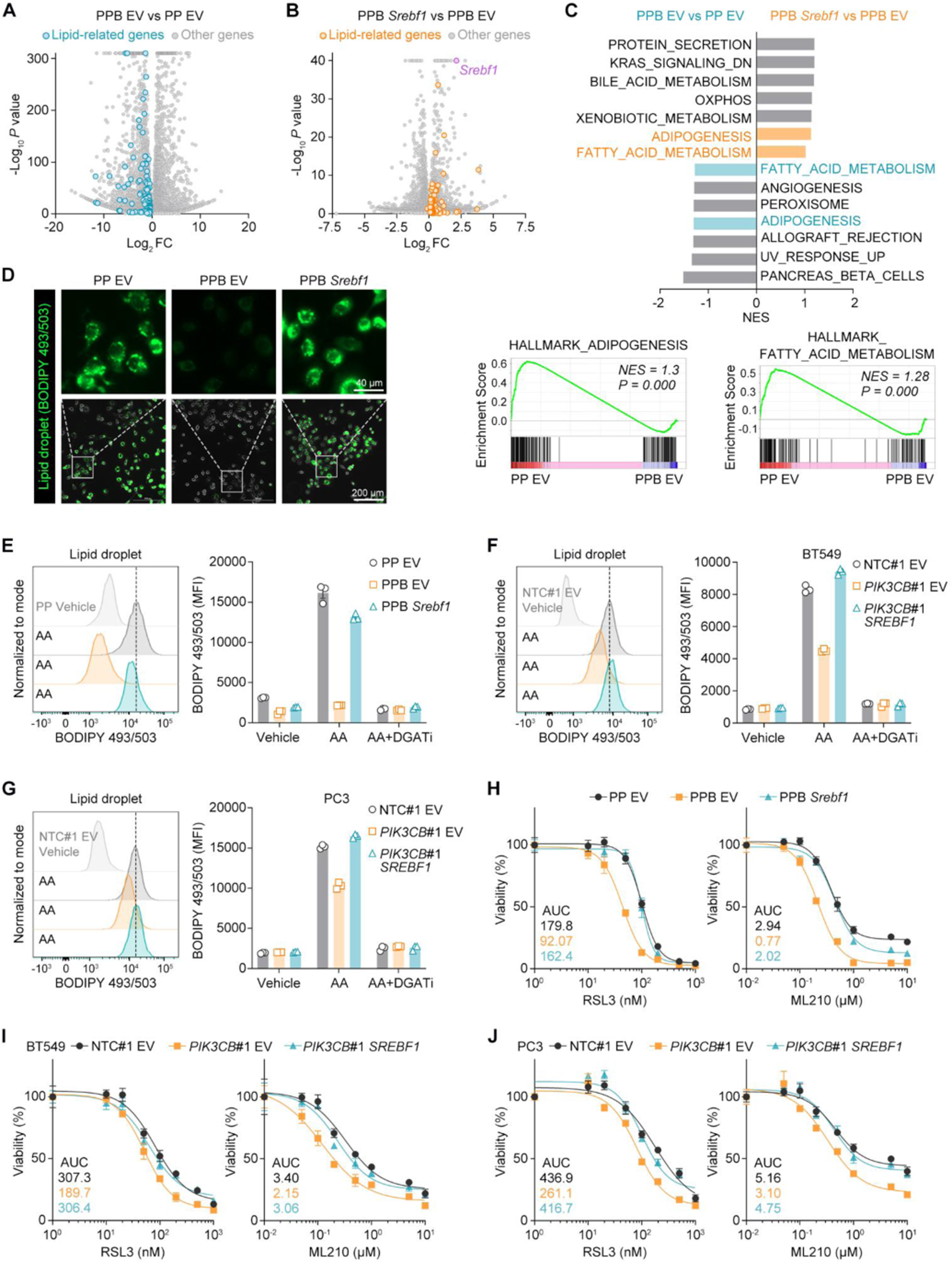
SREBP1 mediates PI3Kβ-dependent lipid metabolic reprogramming, lipid droplet formation, and ferroptosis resistance. **A** and **B,** Volcano plots showing differentially expressed genes from RNA-seq data of PP-EV, PPB-EV, and PPB-Srebf1 cells. Lipid metabolism-related genes are highlighted. **C,** Hallmark pathway enrichment analyses comparing PPB EV versus PP EV cells and PPB Srebf1 versus PPB EV cells. Fatty acid metabolism and adipogenesis pathways are highlighted. Lower panels show representative GSEA plots for adipogenesis and fatty acid metabolism pathways. **D,** Representative BODIPY 493/503 staining images of lipid droplets. Cells were treated with 75 μM AA for 24 h. Scale bars, 200 μm (zoomed-in, 40 μm). **E-G,** Flow cytometry quantification of lipid droplet content. Data are presented as mean ± s.e.m.; n = 3. **H-J,** Dose-response curves of the indicated cells treated with RSL3 or ML210. Data are presented as mean ± s.d.; n = 3.

**Figure S9.**
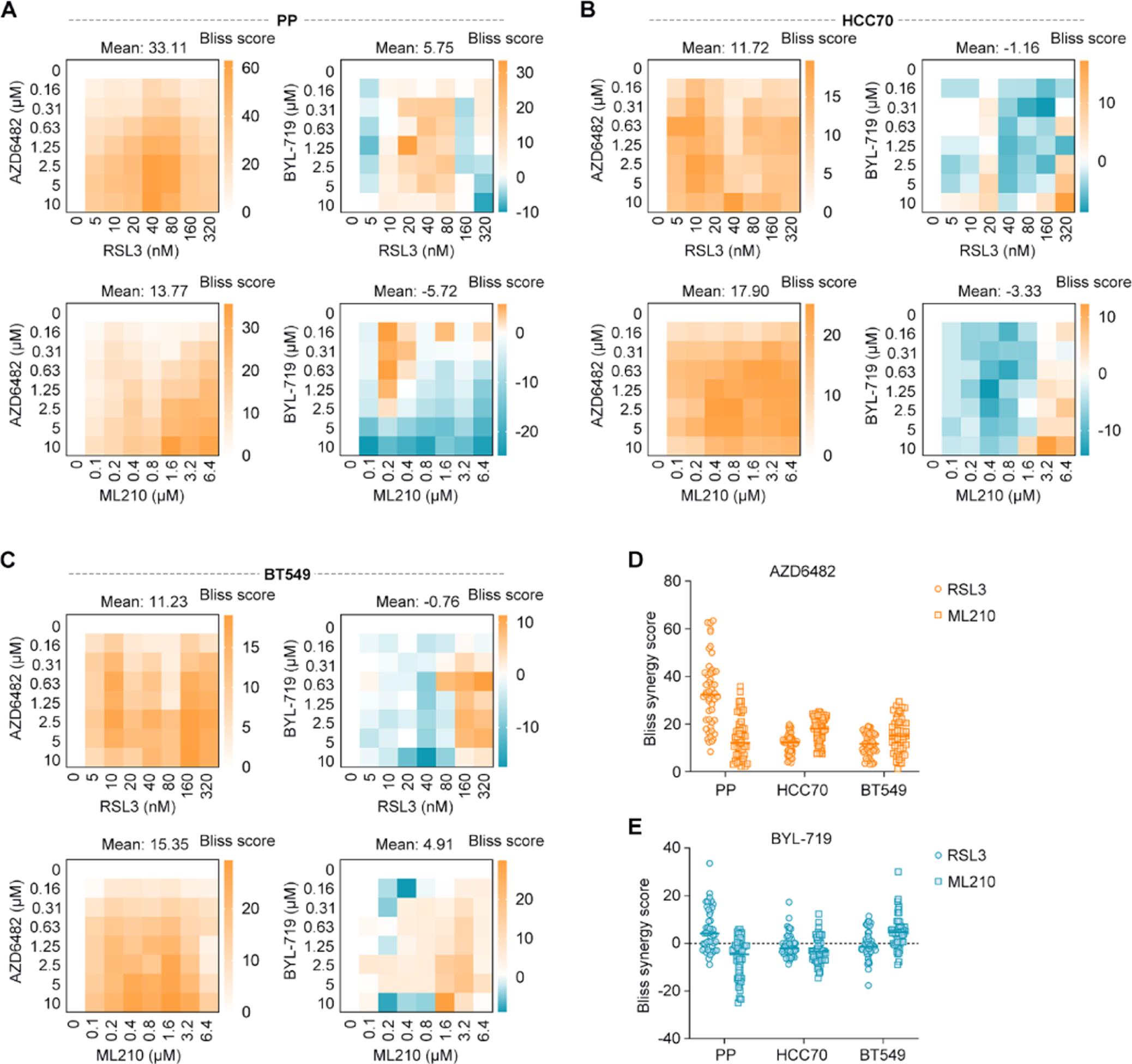
Selective synergy between PI3Kβ inhibition and ferroptosis induction in PTEN-deficient cancer cells. **A-E,** Bliss synergy analysis of AZD6482 or BYL-719 in combination with RSL3 or ML210 in PP, HCC70, and BT549 cells.

**Figure S10.**
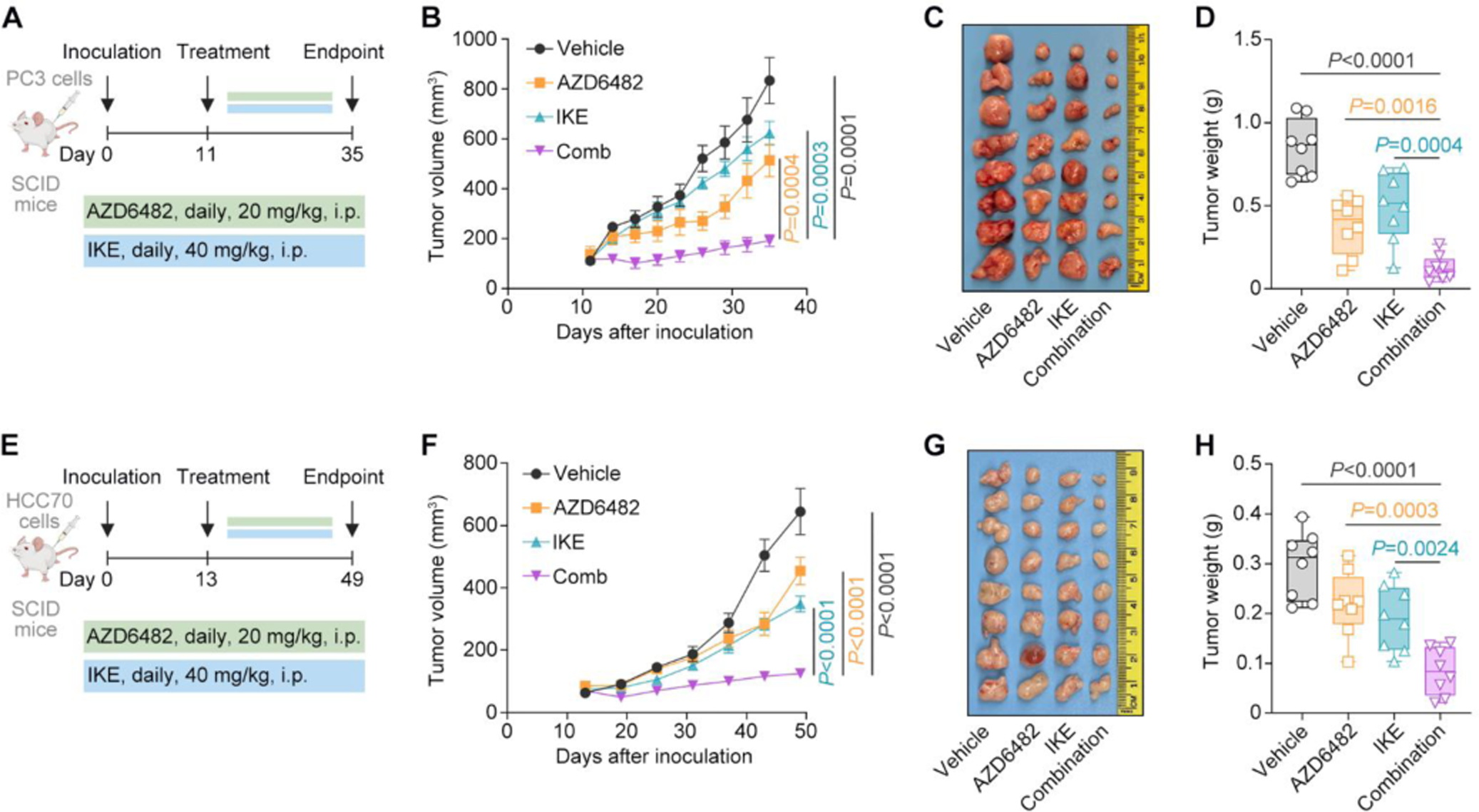
Combined PI3Kβ inhibition and ferroptosis induction suppress PTEN-deficient xenografts growth. **A-D,** Treatment scheme (**A**), tumor growth curves (**B**), representative tumor images (**C**), and tumor weights (**D**) of PC3 xenografts following the indicated treatments. Data are presented as mean ± s.e.m.; two-way ANOVA for tumor volume; unpaired two-tailed t-test for tumor weight; n = 8. **E-H,** Treatment scheme (**E**), tumor growth curves (**F**), representative tumor images (**G**), and tumor weights (**H**) of HCC70 xenografts following the indicated treatments. Data are presented as mean ± s.e.m.; two-way ANOVA for tumor volume; unpaired two-tailed t-test for tumor weight; n = 8.

